# Light-dependent metabolism and proteome allocation in *Synechocystis* sp. PCC 6803: an enzyme-constrained metabolic model across light intensities

**DOI:** 10.64898/2026.07.28.741177

**Authors:** Hossein Firoozabadi, Teddy Groves, Lars Keld Nielsen

**Affiliations:** The Novo Nordisk Foundation Biotechnology Research Institute for the Green Transition, Technical University of Denmark, Lyngby, Denmark; ARC Training Centre for Biopharmaceutical Innovation Australian Institute for Bioengineering and Nanotechnology, The University of Queensland, Brisbane, Queensland, Australia

**Keywords:** Enzyme-constrained metabolic model, Cyanobacteria, Genome-scale metabolic model, Photosynthesis, GECKO, *Synechocystis* sp. PCC 6803

## Abstract

Genome-scale metabolic models (GEMs) are widely used to relate genotype to phenotype and to study how organisms respond to their environment, but conventional flux-balance analysis relies on simplifying assumptions that limit its predictive power. Enzyme-constrained models (ecModels) address this by bounding each reaction by the abundance and turnover number of its catalyzing enzyme. Here we reconstruct ecModels of *Synechocystis* sp. PCC 6803 at light-limited, light-saturated and photoinhibited intensities (27.5, 440 and 1100 µmol photons m⁻² s⁻¹) to study how light shapes its metabolism and enzyme usage. Using the GECKO framework, we constrained the model first by a total protein pool and then by condition-specific quantitative proteomics. The pool-constrained ecModel reproduced the decline in growth at high light as a consequence of a finite proteome, whereas the unconstrained GEM predicted growth to continue rising. Integrating proteomics reduced the median flux variability across reactions by up to two orders of magnitude relative to the conventional GEM. The enzyme budget was dominated by four subsystems (oxidative phosphorylation, transport, photosynthesis and carbon fixation), which together accounted for close to 70% of the minimum enzyme mass in every condition, and the total mass required tracked growth rate rather than light intensity. Enzyme-usage variability analysis found that only about 40% of usages were uniquely determined, a fraction stable across the light gradient. Enzyme constraints thus improve the model’s description of light-dependent cyanobacterial metabolism and identify the functional sectors that carry the metabolic protein budget.

## 1 Background

Genome-scale metabolic models (GEMs) have emerged as valuable tools for investigating the genotype-phenotype relationship[1]. A GEM defines the feasible flux vector space for the organism, and this space can be explored using flux balance analysis (FBA)[2]. Typically, uptake rates of key nutrients are used to constrain the model from which maximum growth rate and theoretical yields can be inferred. However, even when considering a specific cellular objective, e.g., growth, the solution space remains large, which compromises the accuracy of phenotype predictions[3]. Additionally, conventional FBA is unable to capture complex phenomena, such as overflow metabolism, unless specific constraints or objective functions are incorporated.

Enzyme-constrained metabolic models (ecModels), which can be reconstructed through the GECKO framework[4], address the limitations of traditional GEMs. The GECKO framework introduces enzymatic constraints to the metabolic network allowing for a more accurate depiction of cellular metabolism. Notably, recent research[5] demonstrated the potential of ecModels in predicting overflow metabolism in yeast, suggesting that protein limitation may be the underlying reason for this phenomenon. The ecModels, introduced by the GECKO framework, also demonstrated their utility in advancing metabolic engineering applications. An example of this is their role in improving intracellular heme production in *Saccharomyces cerevisiae* [6]. By systematically manipulating gene targets, researchers successfully engineered a yeast strain that achieves a 70-fold increase in heme production compared to the wild type[6].

In this study, we reconstructed enzyme-constrained models of *Synechocystis* sp. PCC 6803, hereafter *Synechocystis*, to study how light intensity shapes its metabolism and enzyme usage across three growth states: light-limited, light-saturated and photoinhibited. We first constrained the model with a total protein pool and investigated how growth responds as light increases, testing whether the enzyme constraint reproduces the decline in growth observed at high light, behaviour that stoichiometry alone does not predict. We then integrated condition-specific quantitative proteomics, which allowed protein usage to be examined across the three light regimes and the resulting proteome allocation to be resolved at the level of metabolic subsystems. Finally, we assessed how each layer of enzymatic constraint, the total protein pool and the measured proteome, shapes the space of feasible flux distributions. Together, these analyses probe how far enzyme constraints extend the description of light-dependent metabolism beyond a conventional GEM.

## 2 Methods

### 2.1 Genome-scale metabolic model curation

A high-quality GEM is the basis for ecModel reconstruction. The genome-scale metabolic model of *Synechocystis* sp. PCC 6803, iJN678 [7], was used as the starting point. The iJN678 model retrieved from the BiGG database [8] differed from the published model in gene count, comprising 622 genes compared with 678 in the original publication [7]. The difference arose from gene naming: the BiGG model used gene names for most reactions whereas the published model used gene loci, so that identical gene names were assigned to genes at distinct loci. A comparative analysis of the two models was conducted to resolve this, and the affected reactions and gene-protein-reaction (GPR) associations were corrected accordingly. After these corrections the GEM comprised 678 genes.

#### 2.1.1 Condition-specific biomass reconstruction

Three light-condition-specific autotrophic biomass reactions were derived from the reference reaction (BIOMASS_Ec_SynAuto) of iJN678 [7], one each for 27.5, 440, and 1100 µmol photons m⁻² s⁻¹ (BIOMASS_Ec_SynAuto_27_5uE, _440uE, _1100uE). Four macromolecular pools for which condition-resolved measurements are available were rescaled to the values reported by Zavřel et al. [9]: total protein, glycogen, chlorophyll a, and total carotenoids (Table S1).

Each pool was rescaled while preserving the internal composition of the reference reaction. The relative amino-acid composition of the protein pool and the relative distribution among carotenoid species were held fixed, and only the group totals were scaled to the measured protein and carotenoid masses. Glycogen and chlorophyll a were set directly from their measured mass using the metabolite molecular weights. Growth-associated maintenance was deliberately not refit per condition: the ATP, water, ADP, proton, phosphate, and pyrophosphate coefficients were kept identical to the reference reaction, so that condition-dependence enters the model only through macromolecular composition. To preserve a consistent biomass basis across conditions, the unmeasured residual precursor pool (nucleotides, lipids, cofactors, and cell-wall components) was rescaled by a single per-condition factor so that total precursor mass, excluding ATP and water, summed to 1 g per mmol of biomass. The residual unmeasured precursor pool required modest upward scaling (factors of 1.10 to 1.16) to close the biomass to 1 g per mmol in all three conditions, well within the tolerance of the procedure and confirming that the measured pools did not distort the biomass mass.

### 2.2 Enzyme-constraint metabolic model (ecModel) reconstruction

The ecModel for *Synechocystis* sp. PCC 6803 was reconstructed from iJN678 [7] using the GECKO 3.2.5 protocol [4]. All reconstruction, curation, and simulation steps were performed in MATLAB R2025b with the RAVEN Toolbox 2.11.1 [10] and the COBRA Toolbox 3.7 [11,12]. Gurobi 13.0.1 (Gurobi Optimization LLC) was used as the optimization solver.

#### 2.2.1 ecModel expansion and initial kcat assignment

The condition-specific GEM was expanded into an ecModel, which introduces each enzyme as a pseudo-metabolite consumed by the reactions it catalyzes and drawing on a shared protein pool, governed by a parameter file (the model adapter) that specifies the biomass reaction, carbon source, taxonomic identifier, and related settings. Enzyme subunit stoichiometry could not be obtained automatically, as *Synechocystis* sp. PCC 6803 is absent from the EBI Complex Portal; subunit counts were therefore curated manually from UniProt [13] and the literature [14–21] and applied to the enzyme-reaction matrix, so that multi-subunit complexes such as the photosystems and RuBisCO carry their correct stoichiometry. Initial turnover numbers were assigned by GECKO’s built-in routines, combining organism-flexible BRENDA matches with DLKcat predictions and selecting one value per reaction. Reactions without a gene-protein-reaction association were assigned a standard turnover number, computed by GECKO as the median of the model’s enzyme-associated Kcat values (20.9 s⁻¹), together with a standard enzyme molecular weight. The protein pool was initialized with the condition-specific total protein content and catalytic protein fraction (Section 2.2.4) and a saturation factor of 0.5. This produced the stage-2 ecModel, which carries untuned turnover numbers and serves as the basis for the evidence-based curation below.

#### 2.2.2 Multi-source kcat evidence base

Beyond the turnover numbers assigned automatically during reconstruction (Section 2.2.1), a dedicated, organism-resolved evidence base was compiled to guide curation. Turnover numbers were compiled from three independent sources for each enzyme-reaction pair in the model: experimentally measured values from BRENDA [22], deep-learning predictions from DLKcat [23], and geometric deep-learning predictions from KcatNet [24].

BRENDA turnover numbers were not assumed to originate from the target organism. Each entry was traced to its source organism, and entries from cyanobacteria, and from *Synechocystis* specifically, were identified; for each EC number the highest wild-type cyanobacterial value was retained in preference to mutant entries.

KcatNet predictions were generated from enzyme sequence and substrate structure. For each reaction, the main non-cofactor substrate was assigned a SMILES string from GECKO’s metabolite dictionary, and the enzyme sequence was taken from the *Synechocystis* reference proteome (UniProt UP000001425). The 786 reactions catalyzed by a single enzyme and carrying both a sequence and a substrate SMILES were submitted to KcatNet (which encodes sequences with the ProtT5 and ESM2 protein language models), run on a GPU. Reactions catalyzed by enzyme complexes were excluded, since their composite identifiers cannot be mapped to a single sequence.

Where several sources gave a value for the same enzyme-reaction pair, a single reference turnover number was selected by an evidence hierarchy that prioritized measured over predicted values and, among predictions, the more recent predictor. In order of preference, this was: an experimental cyanobacterial BRENDA value; the KcatNet prediction; and the DLKcat prediction. KcatNet was preferred over DLKcat as the more recent model, reported to generalize better to enzymes dissimilar from its training data [24]. Pairs for which KcatNet and BRENDA differed by more than tenfold were flagged for manual review.

Agreement between sources was quantified for enzyme–reaction pairs shared by each pair of sources. For every pairing, the median absolute difference in log₁₀ turnover number was computed, together with the fraction of pairs falling within tenfold of one another; the per-reaction correspondence was further assessed by least-squares regression on log₁₀-transformed values, from which the coefficient of determination and slope were obtained.

#### 2.2.3 Kcat curation

Turnover numbers were curated once and applied unchanged across all three light conditions, since turnover numbers are organism-level constants. Curation was performed at light-saturated condition of 440 µmol photons m⁻² s⁻¹, the condition with the highest measured growth rate and therefore the most demanding constraint on enzyme efficiency. Curation proceeded in two phases. In the first, evidence-based phase, the rate-limiting enzymes were identified iteratively, and each was raised to a value drawn from the evidence base (BRENDA, DLKcat, or KcatNet), accepted only when the evidence value exceeded the current model value by at least threefold. Values above an assumed plausibility ceiling of 200 s⁻¹ were rejected as likely over-predictions; the affected enzymes, if still growth-limiting, were handled by the feasibility phase below. In this phase, turnover numbers were set from available evidence, not fitted to the measured growth rate. This phase raised 69 turnover numbers (56 from KcatNet and 13 from DLKcat) but, under the measured protein constraint, brought the model to only 36% of the measured growth rate (0.038 of 0.1044 h⁻¹).

Because evidence-based curation alone left the model well short of the measured growth rate and no further evidence values were available to raise the remaining growth-limiting enzymes, a second feasibility phase used GECKO’s built-in sensitivityTuning function to close the gap. This step adjusted six turnover numbers, each by tenfold: ribulose-bisphosphate carboxylase (RuBisCO, RBPC), photosystem II (PSII), photosystem I (PSI), fructose-bisphosphate aldolase (FBA), fructose-1,6-bisphosphatase (FBP), and thylakoid water transport (H2Otu), bringing the model to the measured growth rate (µ = 0.1044 at 440 µmol photons m⁻² s⁻¹). These six scalings are feasibility adjustments rather than kinetic estimates and are reported separately from the evidence-based raises (Table S2, customKcats_auto.tsv). Because they are not kinetic measurements, they were reverted to their pre-scaling values for the protein-cost analysis, so that the cost ranking reflects evidence-based turnover numbers.

#### 2.2.4 Protein pool, proteomics, and per-condition resolution

Quantitative proteomics for the three light conditions were taken from Zavřel et al. [9], with five biological replicates per condition. In every replicate the summed protein mass fell below the measured total protein content (Ptot), by factors of roughly two to five (Table S3). Each replicate was therefore scaled independently so that its summed mass equaled its measured Ptot. Every protein abundance in replicate *r* was multiplied by the ratio of *Ptot_r* to the summed proteomics mass of *r*, and proteins not detected in a replicate were kept as missing values. The per-replicate Ptot values (Table S4) averaged 402.5, 278.9, and 264.0 mg gDW⁻¹ at 27.5, 440, and 1100 µmol photons m⁻² s⁻¹.

The GECKO protein-pool parameter *f*, the mass fraction of the measured proteome accounted for by enzymes present in the model, was computed per replicate as enzyme-associated protein mass divided by total measured protein mass, using molecular weights from the model. *f* declined monotonically with light intensity, from 0.727 at 27.5 to 0.609 at 1100 µmol photons m⁻² s⁻¹ (Table S4), indicating that a progressively smaller share of total protein resides in metabolic enzymes as light increases. These per-condition values were used in place of GECKO’s default of 0.5. Each ecModel was constrained in two parts: a metabolic constraint that was identical across formulations, and an enzymatic constraint that distinguished them. The metabolic constraint set the exchange-flux bounds per condition: photon uptake capped at the measured value, glucose uptake to zero, bicarbonate uptake left unbounded, and the biomass upper bound to the measured growth rate (Table S5). Note that leaving the bicarbonate exchange unbounded constrains only the uptake of inorganic carbon, not the enzyme catalyzing its membrane transport; the bicarbonate transporter therefore remains subject to the enzyme-concentration constraint like any other enzyme and can appear among the relaxed bottlenecks. Photon uptake rates were provided by T. Zavřel (personal communication), as they were not reported in Zavřel et al. [9] (Table S6).

The enzymatic constraint was implemented in two alternative formulations, both built on the curated Kcat set, which was curated once at 440 µmol photons m⁻² s⁻¹ (Section 2.2.3) and applied unchanged to all conditions. The first was a total protein pool constraint, bounding total enzyme protein by a single pool of size Ptot × f × σ, where σ is the average enzyme saturation factor. Ptot and the per-condition mean f (Table S4) were fixed per condition, and σ was held at the GECKO default of 0.5, no organism-specific value being available for cyanobacteria. The second was a per-enzyme constraint (proteomics-integrated ecModel), fixing each enzyme concentration to its measured absolute abundance. Under these concentrations the maximum attainable growth fell below the measured rate, indicating that the measured abundances of some enzymes were limiting. Enzyme concentrations were therefore relaxed iteratively. At each step the model was solved and the saturated enzymes, those operating at their concentration bound, were identified; among these, the enzyme yielding the largest growth gain upon relaxation was selected, and its concentration bound was raised by the smallest factor on a fixed multiplicative ladder (1.5× to 10□×) that improved growth. Iteration continued until growth reached at least 99% of the measured rate or no bound-limited enzyme yielded further gain. Because turnover numbers were held fixed, reaching the measured rate required large fold-relaxations of some enzyme bounds (up to ∼10³–10□× in the higher-light conditions). Turnover numbers were not modified at this stage: the Kcat curation (Section 2.2.3) and the per-condition concentration relaxation are separate operations, the former adjusting organism-level Kcats and the latter relaxing only per-condition enzyme concentration bounds.

The two enzymatic formulations served different purposes. The total protein-pool formulation was used as the calibration model for turnover-number curation (Section 2.2.3) and for the analyses of the growth response to light and to inorganic carbon (Section 3.4). The three per-enzyme-constrained models, one per light condition, were the basis for the per-condition downstream analyses of enzyme allocation, capacity usage, and proteome composition across the light gradient. For each condition, the minimum protein-pool usage was obtained by a two-step optimization on the resolved model: growth was first maximized, then the total enzyme mass (the protein-pool exchange flux) was minimized subject to growth remaining at ≥99% of that maximum.

#### 2.2.5 Flux and enzyme-usage variability analysis

Flux variability analysis (FVA) was used to quantify how the enzyme constraints shaped the space of feasible solutions, in two complementary ways. Both analyses used Gurobi and were performed per light condition on the same models described above, with growth fixed at 99% of each model’s maximum so that all formulations were compared at a common growth rate.

The first analysis measured the size of the feasible flux space. For each light condition, FVA was run on three formulations, the conventional GEM, the total protein-pool ecModel, and the proteomics-integrated ecModel, and the minimum and maximum flux of every reaction was recorded. The variability range of each reaction was taken as the difference between its maximum and minimum flux, and the distribution of these ranges was summarized by the median over reactions able to vary (range > 10⁻□ mmol gDW⁻¹ h⁻¹). Reactions were further grouped into four variability classes by their range: fixed (< 10⁻□), tightly constrained (10⁻□–10⁻²), moderate (10⁻²–1), and broadly variable (> 1 mmol gDW⁻¹ h⁻¹).

The second analysis assessed how uniquely the enzyme usage itself was determined. FVA was applied to the enzyme-usage variables of each condition-specific proteomics-integrated model, again at fixed growth, recording the range each enzyme’s usage could take across alternative optimal solutions. An enzyme was classed as forced, that is uniquely determined, when its usage range was below 10⁻□ mg gDW⁻¹ or below 5% of its mean usage, and as discretionary otherwise.

## 3 Results

### 3.1 A sparse measured Kcat core motivates a multi-source evidence stack

Reconstructing an enzyme-constrained model requires a turnover number for every enzyme–reaction pair, yet measured Kcat values are scarce even for the most intensively studied organisms. For Escherichia coli, the biochemically best-characterized organism, in vitro Kcat is known for only about 10% of enzyme-catalysed reactions [25], and even in yeast, where enzyme-constrained models are most mature, conventional BRENDA/SABIO-RK matching constrains only ∼60% of enzyme-annotated reactions, rising to ∼90% only once sequence-based prediction is added [26]. If the kinetic record is this incomplete for the model organisms, it is far sparser for a non-model phototroph. For *Synechocystis* sp. PCC 6803 the organism-specific gap is severe: of the 1152 enzyme–reaction pairs in the model, only 54 (4.7%) could be assigned a turnover number measured in *Synechocystis* sp. PCC 6803 itself, and only 75 (6.5%) a value measured in any cyanobacterium (Fig. 1A), an order of magnitude below the coverage that organism-matched databases provide for E. coli or yeast. The measured cyanobacterial core is therefore far too small to parameterize the model on its own. Coverage could be extended only by relaxing organism specificity: cross-organism BRENDA matching reached 663 pairs (57.6%), but at the cost of drawing predominantly on distantly related taxa, while the sequence- and structure-based predictors gave the broadest reach, DLKcat 851 pairs (73.9%) and KcatNet 786 (68.2%) (Fig. 1A). Even with all three sources combined, 89 pairs (7.7%) carried no evidence and retained the model’s standard turnover number.

**Fig 1.**
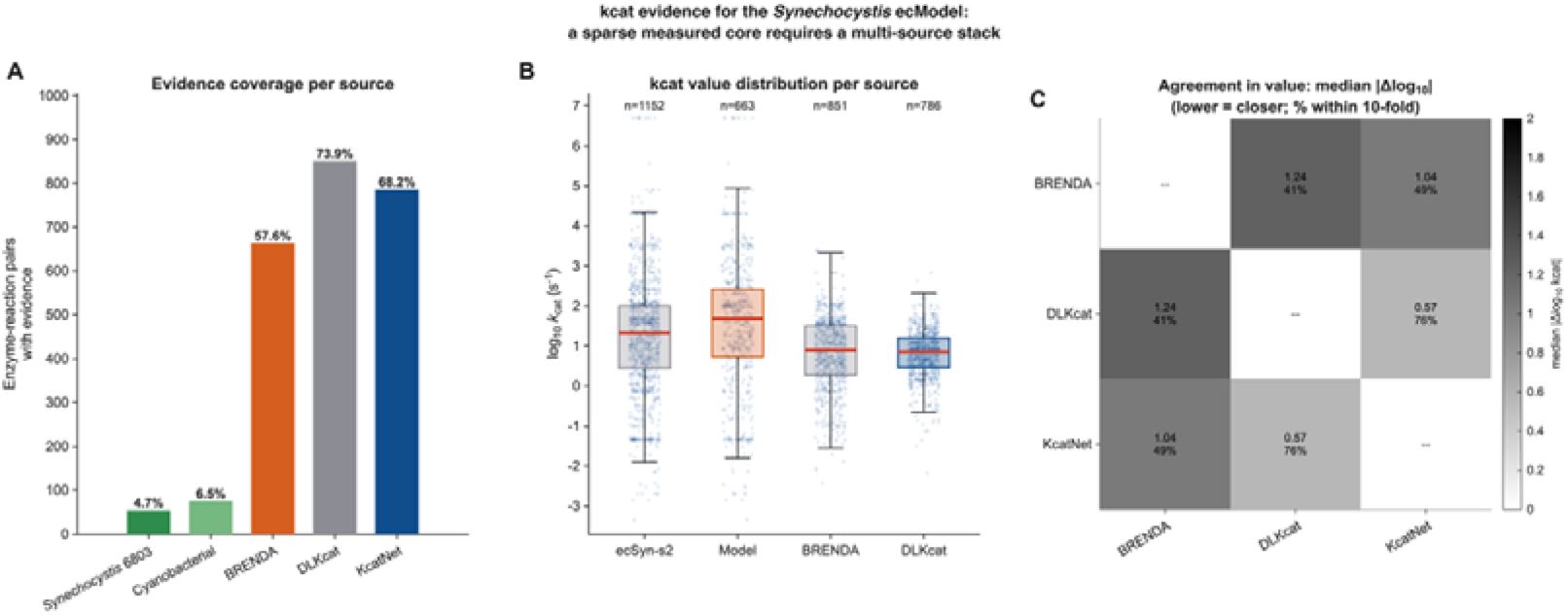
Kcat evidence for the *Synechocystis* sp. PCC 6803 ecModel. Kcat values for the 1152 enzyme–reaction pairs were compiled from BRENDA (organism-resolved), DLKcat, and KcatNet. (A) Coverage per source: number of enzyme–reaction pairs with a value and the percentage of all 1152 pairs. (B) Distribution of log_₁₀_ Kcat per source (box: median and IQR; whiskers: 1.5× IQR; points: individual pairs; n above each). (C) Pairwise agreement for overlapping pairs: median absolute difference in log_₁₀_ Kcat (top value) and percentage of pairs within tenfold (bottom value); greyscale encodes median |Δlog_₁₀_|.

### 3.2 The Kcat sources disagree at the level of individual reactions

The three sources have broadly overlapping value distributions with comparable central tendencies (Fig. 1B), yet they agree poorly when compared reaction by reaction (Fig. 1C). Across the three sources, the median absolute log₁₀ difference for overlapping pairs ranged from 0.57 to 1.24, and only 41 to 76% of pairs fell within tenfold of one another (Fig. 1C). Agreement was weakest between BRENDA and DLKcat (median |Δlog₁₀| = 1.24; 41% within tenfold). The closest agreement was between DLKcat and KcatNet (0.57; 76% within tenfold). Because both are sequence-based predictors trained on overlapping experimental data, however, their mutual concordance reflects shared method and training signal rather than independent corroboration. Pairwise regression confirmed the absence of per-reaction correlation. Coefficients of determination were uniformly low (r² = 0.017 to 0.075) and all slopes lay far below the identity line (Fig. S1), indicating that the predictors compress the dynamic range of measured values rather than reproducing it. This per-reaction dispersion is in line with reports that experimentally measured Kcat data are sparse and noisy [26] and that DLKcat loses predictive accuracy for enzymes dissimilar to its training data [27].

Taken together, the coverage and agreement analyses establish that the model cannot be parameterized by adopting any single source wholesale. Instead, the three sources were reconciled through an evidence hierarchy that prioritizes organism-proximal measured values over inferred and predicted ones (Section 2.2.2), and turnover-number curation was restricted to the enzymes that limit growth rather than applied indiscriminately across the network (Section 3.4).

### 3.3 Enzyme constraints expand the metabolic network and couple it to the proteome

Enzyme-constraint reconstruction expanded iJN678 (866 reactions, 795 metabolites) into eciJN678, comprising 2013 reactions, 1470 metabolites, and 678 enzymes. Two enzymatic formulations were built on this scaffold (Section 2.2.4): a total protein-pool model and a proteomics-integrated model in which enzymes are fixed to their measured abundances. Across the three light conditions, roughly 70% of the enzymes (468–480) carried a measured abundance. Quantitative proteomics recovered only part of the total protein content, with summed detected mass falling two- to fivefold below the measured Ptot across replicates (Table S3), consistent with incomplete recovery before quantification. The catalytic protein fraction *f*, the share of total protein residing in model enzymes, declined with light intensity, from 0.727 at 27.5 to 0.609 at 1100 µmol photons m⁻² s⁻¹ (Table S4), indicating that a progressively smaller fraction of the proteome is invested in metabolic enzymes as light increases.

**Fig. S1.**
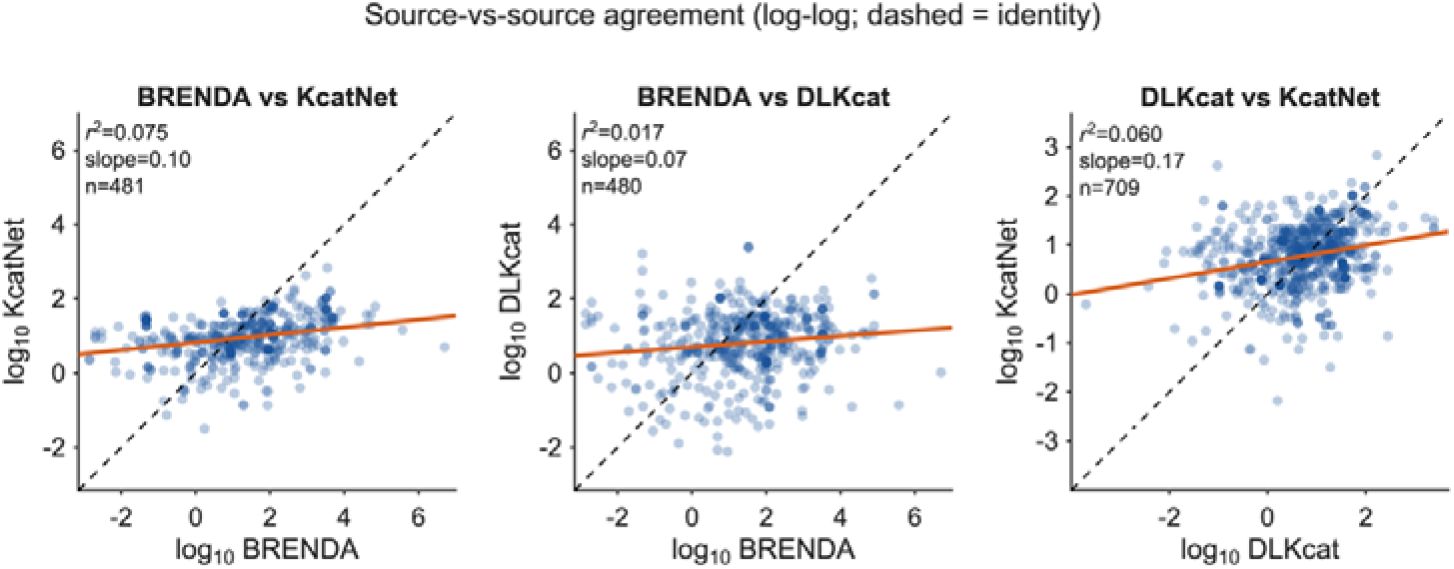
Source-versus-source agreement of Kcat values at the per-reaction level. Pairwise log_₁₀_ Kcat comparison for enzyme–reaction pairs shared by both sources. Each point is one pair; dashed line, identity (y = x); solid line, least-squares fit. r², slope, and n are given in each panel.

### 3.4 Enzyme constraints reproduce the high-light growth decline under a total protein-pool limit

A conventional GEM cannot reproduce the decline in cyanobacterial growth at high light. Without enzymatic constraints, iJN678 predicts growth increasing linearly with photon uptake without bound (Fig. 2A) and a constant biomass yield fixed at the stoichiometric ceiling (2.03 × 10⁻³ h⁻¹ per mmol photon; Fig. 2B). This monotonic response contradicts the established physiology of *Synechocystis* sp. PCC 6803, in which growth increases with light up to a saturating optimum and then declines at supra-optimal intensities through photoinhibition [28].

**Fig. 2.**
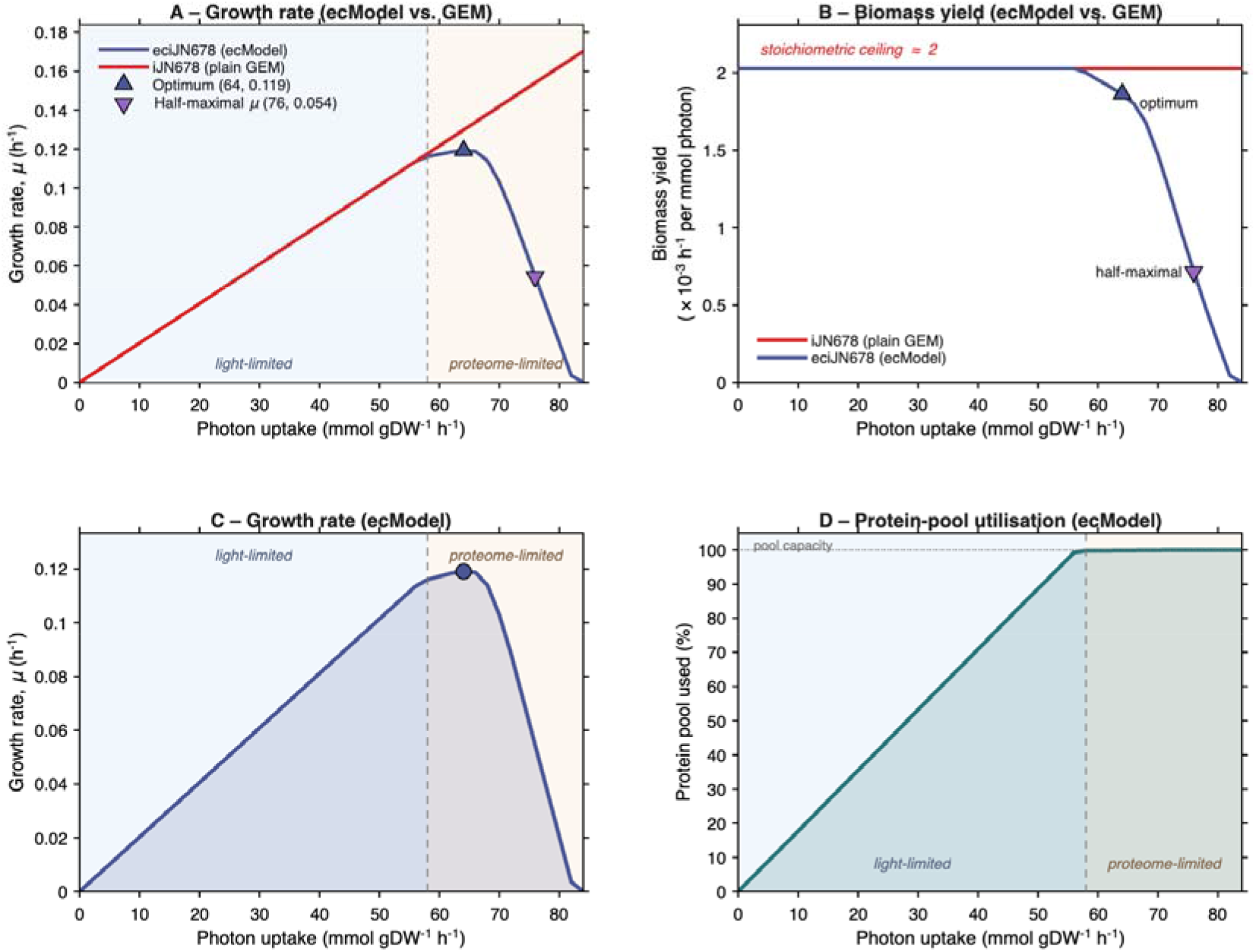
Enzyme constraints reproduce the high-light growth decline under a total protein-pool limit. Growth simulation over a photon-uptake series, for the enzyme-constrained model (eciJN678, navy) and its parent GEM (iJN678, red). (A) Growth rate versus photon uptake. (B) Biomass yield per photon. (C) ecModel growth rate and (D) minimum protein-pool utilisation (fraction of the enzyme pool needed to sustain maximum growth) versus photon uptake. Dashed line: onset of proteome limitation (first photon uptake where ecModel growth falls below the GEM, at 58 mmol gDW⁻¹ h⁻¹). Shading separates the light-limited (ec = GEM) and proteome-limited (ec < GEM) regimes; ▴ (A) and • (C) mark the growth optimum, and ⍰ (A) the half-maximal point.

Imposing a total protein-pool constraint recovered this phenotype. In eciJN678, growth increased with photon uptake up to an optimum at 64 mmol gDW⁻¹ h⁻¹ (µ = 0.119 h⁻¹), then declined monotonically, reaching zero near 84 mmol gDW⁻¹ h⁻¹ (Fig. 2A,C). Biomass yield behaved likewise: it held at the stoichiometric ceiling while the ecModel tracked the GEM, then fell away once the pool became binding, collapsing to zero by ∼84 mmol (Fig. 2B). This response reflects the finite protein pool. At low photon uptake growth was light-limited: the enzyme-constrained solution was identical to the unconstrained GEM and the pool retained usable spare capacity (Fig. 2A,D). Proteome limitation set in where the two models diverged, at photon uptake of 58 mmol gDW⁻¹ h⁻¹, the point at which the protein pool became binding and ecModel growth dropped below the GEM (Fig. 2A).

Critically, this onset is distinct from the growth optimum. From photon uptake of 58 mmol gDW⁻¹ h⁻¹, the pool held growth below the unconstrained ideal, yet growth continued to rise to its maximum at photon uptake of 64 mmol gDW⁻¹ h⁻¹ because light remained co-limiting over this short interval; across it the pool stayed pinned at capacity (99.86–99.91%; Fig. 2D) rather than filling further. The optimum photon uptake at 64 mmol gDW⁻¹ h⁻¹ is therefore the peak of the proteome-limited branch, not its onset. Beyond the optimum the ecModel declined while the GEM kept rising over the same range (Fig. 2A); as the two differ only in the pool constraint, the decline is attributable to the finite proteome rather than to reaction stoichiometry. The growth optimum is thus set by proteome capacity.

This is the phototrophic counterpart of overflow metabolism, which enzyme-constrained genome-scale models reproduce in *S. cerevisiae*: the respiration-to-fermentation switch (the Crabtree effect) emerges from the finite enzyme pool rather than from network stoichiometry, a behaviour conventional GEMs cannot capture [29].

The same constraint governed the joint dependence of growth on light and inorganic carbon (Fig. 3). Across the photon × bicarbonate phase plane, the unconstrained GEM predicted growth increasing monotonically toward the high-light, high-carbon corner (Fig. 3B), whereas eciJN678 confined growth to a bounded interior region with an optimum at intermediate photon uptake (Fig. 3A), beyond which additional light reduced growth. Enzymatic constraints therefore bound the feasible growth region along both the light and carbon uptakes, converting the open-ended response of the conventional GEM into a bounded, finite optimum.

**Fig. 3.**
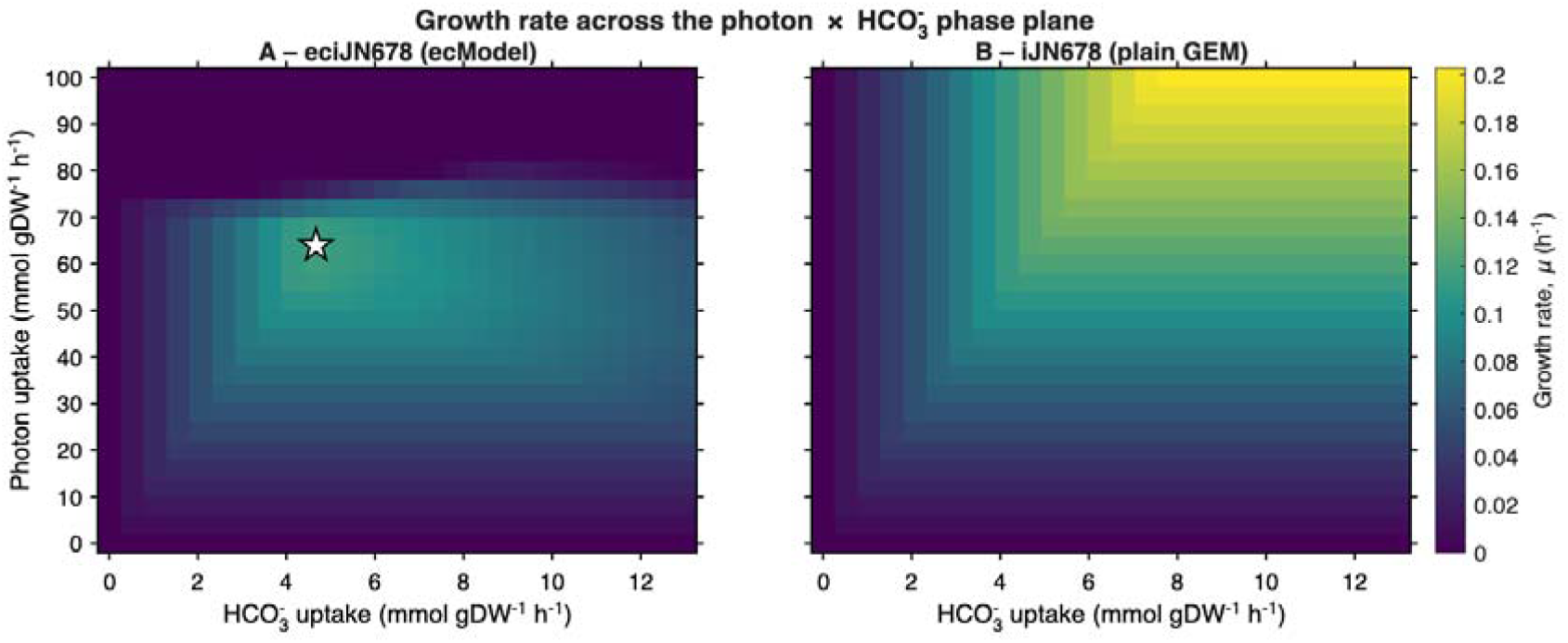
Growth across the photon × bicarbonate uptake phase plane. Growth rate (µ, colour) for (A) the enzyme-constrained model (eciJN678) and (B) its parent GEM (iJN678), on a shared colour scale. White star in (A): global growth optimum.

### 3.5 Proteomics-integrated ecModel

#### 3.5.1 Measured enzyme abundances under-support growth, requiring targeted enzyme relaxation

In the second formulation, each enzyme was fixed to its measured abundance rather than drawing on a shared pool, giving a condition-specific model for every light intensity (Section 2.2.4). Because quantitative proteomics did not cover the entire enzyme complement, the enzymes fall into three groups by the origin of their concentration bound: those set directly from a measured abundance, those whose measured bound was relaxed to achieve growth (Section 2.2.4), and those without a measured value, which retain the model’s default bound. Table 1 reports the number of enzymes in each group and their contribution to the minimum enzyme mass in each light condition.

**Table 1.**
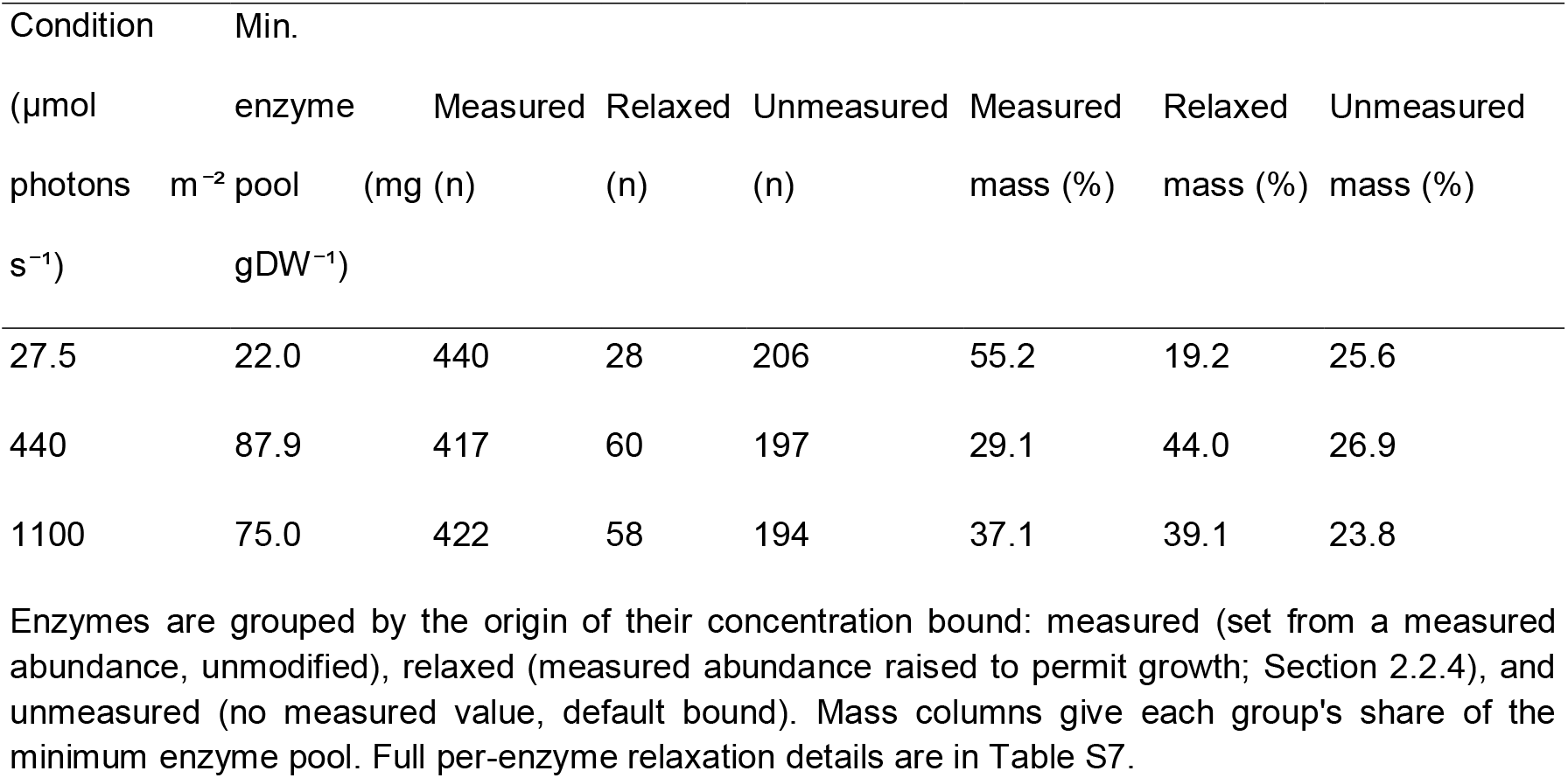
Enzyme composition of the proteomics-integrated model, by the origin of each enzyme’s concentration bound, for the three light conditions. Enzymes are grouped by the origin of their concentration bound: measured (set from a measured abundance, unmodified), relaxed (measured abundance raised to permit growth; Section 2.2.4), and unmeasured (no measured value, default bound). Mass columns give each group’s share of the minimum enzyme pool. Full per-enzyme relaxation details are in Table S7.

Fixing each enzyme to its measured abundance is a considerably stricter constraint than the shared protein pool, as it removes the freedom to redistribute enzyme mass between reactions. Under the measured abundances alone, the attainable growth rate fell far below the experimental value in all three light conditions, to 9.6%, 1.4%, and 1.5% of the measured rate at 27.5, 440, and 1100 µmol photons m⁻² s⁻¹, respectively, indicating that a subset of enzymes was present at concentrations too low to support the observed growth. Relaxing these growth-limiting bounds (Section 2.2.4) recovered essentially the measured growth rate in every condition, reaching 99.4%, 100%, and 99.3% of the experimental value, and required 28, 60, and 58 enzyme bounds to be relaxed, respectively (Table S7). Turnover numbers were not re-tuned at this stage, having already been curated on the total protein-pool ecModel (Section 2.2.3); the per-condition step adjusted only enzyme concentration bounds. Because these relaxations act on enzymes whose measured abundances were often very low, a large fold-change corresponds to only a small absolute change, and the relaxed bounds remain comparable to the default bound applied to enzymes without a measured value (Table S7). They are, moreover, only upper limits rather than usages: when protein usage is minimized, the enzyme mass carried lies far below them, so the relaxations mark where the measured abundances were most limiting under the fixed turnover numbers, not the enzyme quantities the model ultimately uses.

By mass, directly measured enzymes accounted for the largest share of the minimum enzyme pool at the light-limited condition (55.2%) and a smaller share at the light-saturated and photoinhibited conditions (29.1% and 37.1%), where relaxed bounds contributed more (44.0% and 39.1%); enzymes without a measured abundance contributed a comparable fraction throughout (23.8–26.9%; Table 1). The measured share therefore fell, and the relaxed share rose, as growth increased, consistent with more enzymes reaching their measured limit at higher growth.

We next determined the minimum protein-pool usage for each light condition, defined as the smallest enzyme mass able to sustain that condition’s growth rate. Protein usage was highest at the light-saturated condition (87.9 mg gDW⁻¹ at 440 µmol photons m⁻² s⁻¹), which also reached the highest growth rate, followed by the photoinhibited (75.0 mg gDW⁻¹, 1100 µmol photons m⁻² s⁻¹) and lowest at the light-limited (22.0 mg gDW⁻¹, 27.5 µmol photons m⁻² s⁻¹) condition. Protein usage therefore matched the growth rates, not the light intensities, with faster growth requiring a larger enzyme mass, consistent with the finite-proteome behavior seen under the total protein-pool constraint.

#### 3.5.2 Saturation is confined to a small subset of enzymes

To assess how heavily the resolved proteome is used, we computed each enzyme’s capacity usage at the minimum-pool solution, defined as its usage divided by its concentration bound (the fraction of the available enzyme). Enzymes without a measured abundance were excluded, as their bound is the default value of 1000 mg gDW⁻¹ assigned by the GECKO pipeline rather than a measurement; the analysis therefore covers the 468–480 enzymes carrying a measured abundance, those set directly from proteomics and those whose bound was relaxed (Table 1) in each condition.

In all three light conditions the distribution was strongly skewed toward low usage (Fig. 4): most enzymes operated far below their bound, with a median capacity usage of only 0.2%, 1.7%, and 1.2% at 27.5, 440, and 1100 µmol photons m⁻² s⁻¹, and 69–82% of enzymes using less than one-tenth of their capacity. The measured proteome therefore carries substantial spare catalytic capacity, consistent with reports that a large fraction of the *Synechocystis* carbon- and light-assimilation proteome is not fully utilized [30]. Only a small subset ran near saturation (capacity usage ≥ 0.9): 11, 27, and 21 enzymes across the three conditions, representing 2.4%, 5.7%, and 4.4% of the measured set.

**Fig. 4.**
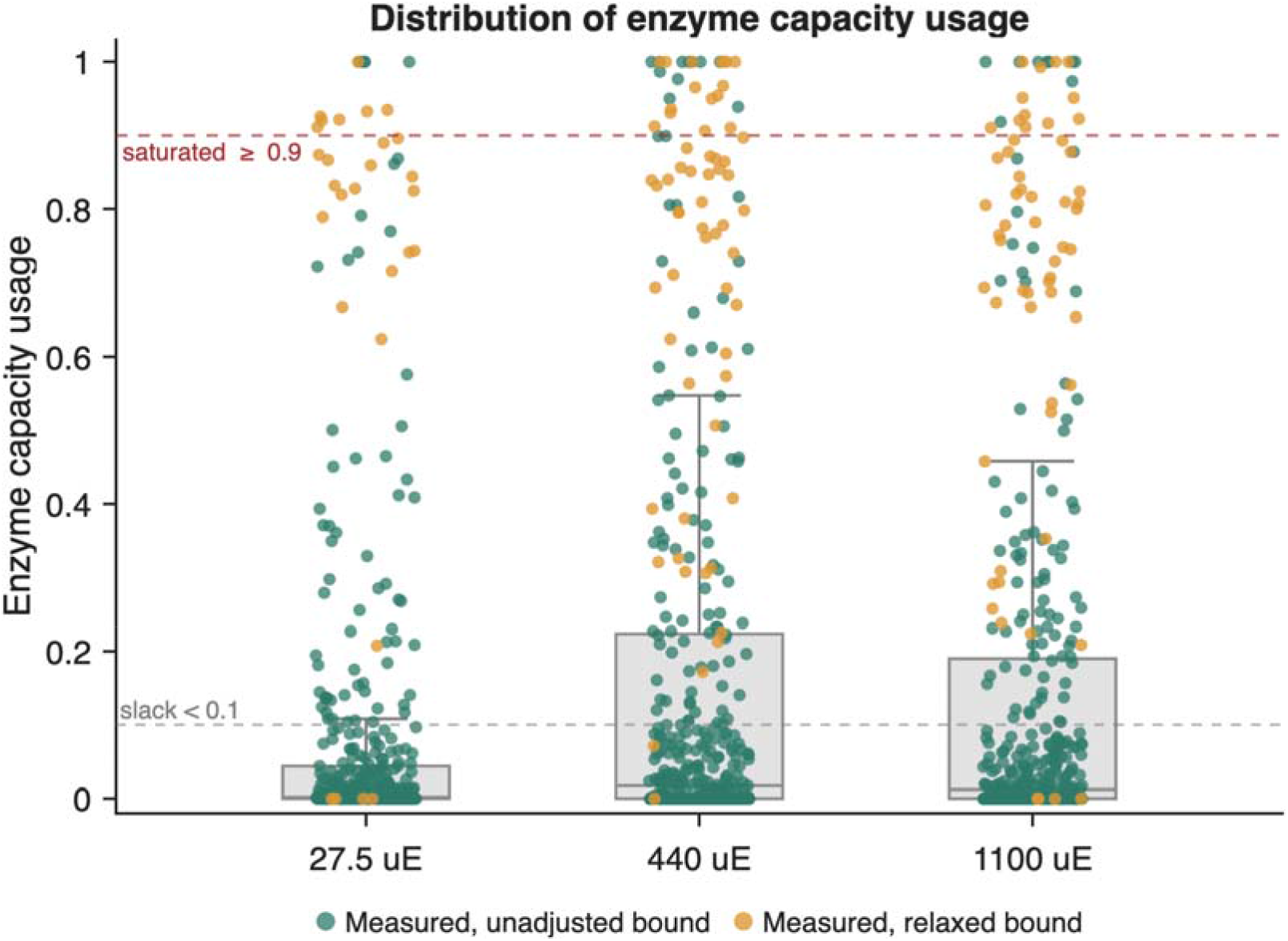
Distribution of enzyme capacity usage across the three light conditions. Capacity usage (enzyme usage ÷ concentration bound, at the minimum-pool solution) for every enzyme with a measured abundance, at 27.5, 440, and 1100 µmol photons m⁻² s⁻¹. Each point is one enzyme: green, bound set from the measured abundance; amber, bound relaxed to permit growth (capacity expressed relative to the relaxed bound). Boxes show median and interquartile range; dashed lines mark the saturation (0.9) and slack (0.1) thresholds. Enzymes without a measured abundance are excluded.

Both the median usage and the number of saturated enzymes were highest at 440 µmol photons m⁻² s⁻¹, the fastest-growing condition, so saturation followed growth rate rather than incident light. Even at the fastest growth, only a small fraction of enzymes approached their catalytic limit, therefore tracked the growth rate rather than the incident light, and even at the fastest growth only a small fraction of the proteome approached its catalytic limit, with the majority held in reserve.

#### 3.5.3 Proteome allocation concentrates in a few major subsystems and scales with growth rate

To examine how the enzyme budget is distributed among metabolic functions, we aggregated the minimum-pool enzyme usage by subsystem, assigning each enzyme’s usage to the subsystems of the reactions it catalyzes in proportion to the flux each reaction carried. Aggregation to the subsystem level is deliberately robust: interchange between isozymes or alternative electron carriers, which leaves an individual reaction’s usage ambiguous, cancels within a subsystem total. Because the three condition-specific models differ in their measured proteome, total protein content and growth rate, that is, they represent three distinct acclimation states rather than one system observed repeatedly. Thus, we report the enzyme mass each state commits in absolute terms (mg gDW⁻¹) rather than as a normalized fraction, so that the differences in total budget and in growth are retained rather than divided out.

The enzyme budget was dominated by the same four subsystems in every condition (Fig. 5). Oxidative phosphorylation, transport, photosynthesis, and carbon fixation together accounted for close to 70% of the minimum enzyme mass at each light intensity, while individual biosynthetic pathways each contributed a few percent or less. This concentration of the proteome into the energy-conserving, light-harvesting and carbon-assimilating machinery, together with its scaling with growth rate, is consistent with quantitative proteomics of *Synechocystis*, in which these functional sectors dominate the proteome mass and change approximately linearly with growth rate [30].

**Fig. 5.**
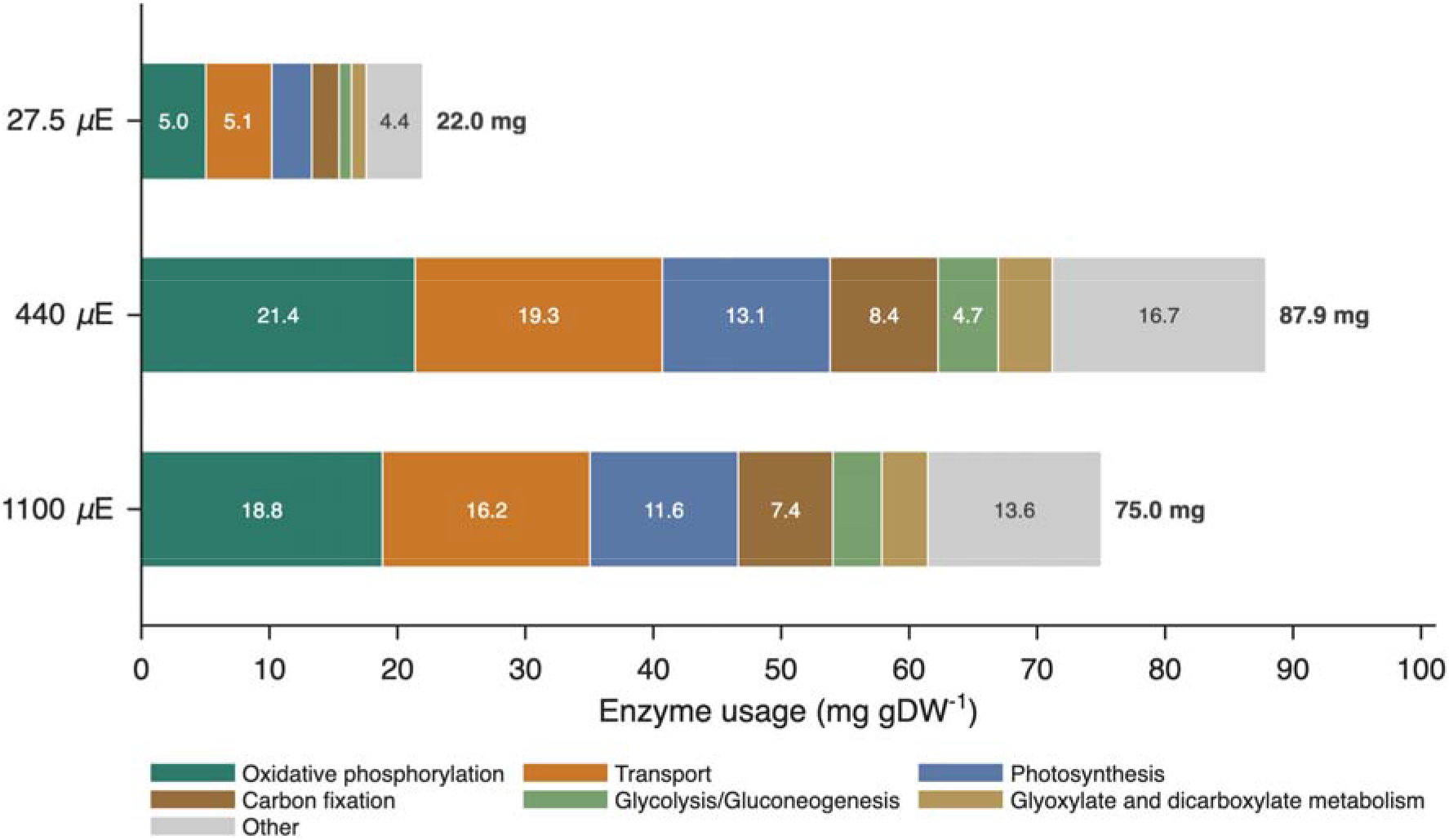
Enzyme mass allocated to metabolic subsystems in each light condition. Minimum-pool enzyme usage (mg gDW⁻¹) partitioned by subsystem at 27.5, 440, and 1100 µmol photons m⁻² s⁻¹, with each enzyme’s usage assigned to the subsystems of the reactions it catalyzes in proportion to their flux. Bar length is the total minimum enzyme pool of that condition (labelled at right); segments give the mass of the six highest-usage subsystems, with all remaining subsystems combined as “Other”. Growth was fixed at each model’s maximum achievable rate (99% of the measured value). Values are the minimum enzyme allocation required by the ecModel.

The total enzyme mass required tracked the growth rate. It was largest at the light-saturated condition (87.9 mg gDW⁻¹ at 440 µmol photons m⁻² s⁻¹), which reached the highest growth rate, and smaller at both the photoinhibited (75.0 mg gDW⁻¹, 1100 µmol photons m⁻² s⁻¹) and light-limited (22.0 mg gDW⁻¹, 27.5 µmol photons m⁻² s⁻¹) conditions. Each dominant subsystem scaled in the same direction: from the light-limited to the light-saturated condition, oxidative phosphorylation rose from 5.0 to 21.4, photosynthesis from 3.1 to 13.1, carbon fixation from 2.1 to 8.4, and transport from 5.1 to 19.3 mg gDW⁻¹, all contracting again at the photoinhibited condition. The cell therefore met faster growth by scaling the whole enzyme investment up and down with growth rate, while keeping the broad division among functional sectors approximately constant, rather than by qualitatively re-partitioning the proteome between conditions. This differs from the trade-off described by bacterial growth laws, in which the ribosomal proteome fraction expands with growth rate at the expense of metabolic and biosynthetic sectors [31]; the enzyme-constrained model resolves only the metabolic proteome and does not represent the ribosomal sector that drives that trade-off, so the metabolic sectors here scale together with growth rather than being displaced.

Two features of this analysis should be read with care. First, the masses are the minimum enzyme allocation the model needs to sustain each condition’s growth, not the regulated proteome the cell expresses; they describe the metabolic demand implied by the flux state, and the near-constant sector division partly reflects that flux through the biomass-coupled core scales with growth under fixed turnover numbers. Second, part of the transport mass is carried by transporters without a measured abundance (for example the sodium/proton antiporter and thylakoid water transport), whose concentration bounds take the model default (Table 1); the transport totals are therefore more bound-dependent than those of the energy and carbon-fixation sectors, whose principal enzymes carry measured or relaxed abundances. For this reason, we restrict the interpretation to the subsystem level, where these individual assignments are aggregated, and do not draw conclusions from individual transporter usages.

### 3.6 Flux and enzyme-usage variability

Constraining a metabolic model with enzyme information narrows the set of flux distributions it admits, but it does not necessarily pin down a unique solution. We used flux variability analysis (FVA) to characterize both effects: first, how much the enzyme and proteomics constraints shrink the feasible flux space relative to the genome-scale model (section 3.6.1); and second, how much of the resulting per-enzyme protein allocation is uniquely determined rather than free to vary across equally-optimal solutions (section 3.6.2).

#### 3.6.1 Enzyme constraints reduce flux variability

A recognized advantage of enzyme-constrained models is that they shrink the space of flux distributions relative to the underlying genome-scale model, because each reaction now competes for a finite protein budget [29]. To quantify this for the *Synechocystis* ecModel, we performed flux variability analysis (FVA) at all three light intensities on three model formulations: the conventional GEM, the ecModel constrained only by the total protein pool, and the proteomics-integrated ecModel with per-enzyme abundance bounds. Within each condition all three formulations were held at the same growth rate, so differences in variability reflect the enzyme constraints alone.

Each layer of constraint contracted the feasible flux space (Fig. 6). Relative to the GEM, the protein-pool constraint reduced the median flux-variability range across reactions that could vary by roughly one to two orders of magnitude at every condition, and the proteomics-integrated model reduced it further at the two lower light intensities. At the light-saturated condition (440 µmol photons m⁻² s⁻¹) the median range fell from 5.3 × 10⁻² mmol gDW⁻¹ h⁻¹ in the GEM to 4.0 × 10⁻³ under the total protein pool and 5.4 × 10⁻□ with proteomics integration, a 97-fold reduction; the corresponding reductions were 34-fold at 27.5 and 27-fold at 1100 µmol photons m⁻² s⁻¹. The distribution of per-reaction ranges shifted accordingly (Fig. 6): at 440 µmol photons m⁻² s⁻¹ the fraction of essentially fixed reactions (variability below 10⁻□) rose from 27% in the GEM to 42% under the pool constraint and 53% with proteomics, while the fraction of broadly variable reactions (range above 1) fell from 28% to 14% to 4%, with the same trend at the other intensities. Adding the measured proteome therefore not only tightened the typical reaction but also removed most of the widely variable ones. The reduction relative to the genome-scale model is consistent with enzyme-constrained models of other organisms [29,32–34].

**Fig. 6.**
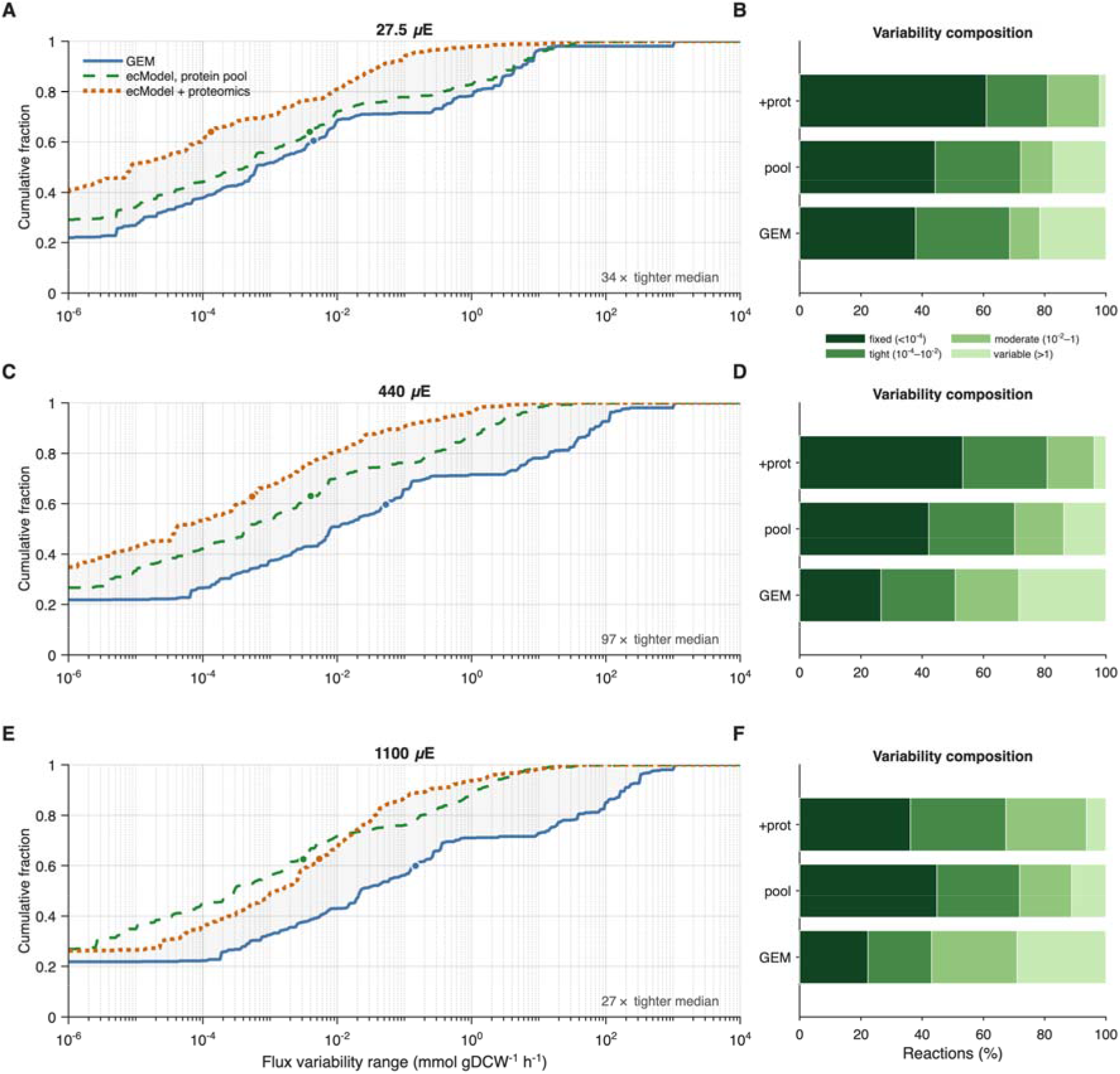
Enzyme constraints reduce the feasible flux space across the light gradient. Flux variability analysis of three model formulations — the conventional genome-scale model (GEM), the ecModel constrained by the total protein pool, and the proteomics-integrated ecModel — at 27.5 (**A, B**), 440 (**C, D**) and 1100 (**E, F**) µmol photons m⁻² s⁻¹. Within each condition all three formulations were held at the same growth rate. **(A, C, E)** Cumulative distribution of the per-reaction flux-variability range (max − min flux); curves further to the left indicate a more constrained network. Circles mark the median over reactions that can vary (range > 10⁻□); the shaded band spans the GEM and proteomics-integrated curves (the solution space removed by the constraints), and the fold reduction in median range from the GEM to the proteomics-integrated model is annotated. **(B, D, F)** Composition of the reaction set by variability class — fixed (< 10⁻□), tightly constrained (10⁻□–10⁻²), moderate (10⁻²–1), and broadly variable (> 1 mmol gDW⁻¹ h⁻¹). At 1100 µmol photons m⁻² s⁻¹ the proteomics-integrated model was infeasible at the operating growth rate and was evaluated with its growth lower bound relaxed to zero; its median range therefore exceeds that of the protein-pool model at this condition, unlike at the two lower intensities.

#### 3.6.2 Enzyme usage is not uniquely determined

A single optimal solution does not fix each enzyme’s usage uniquely, where flux can route through alternative isozymes or carriers, individual usages vary across equally optimal solutions while the total pool stays fixed. To quantify how much of the per-enzyme allocation is genuinely determined, we ran flux variability analysis on the enzyme-usage variables of each condition-specific model, holding growth at its maximum, and recorded the range (maximum − minimum usage) each enzyme could take across alternative optima. Enzymes whose range was negligible (below 10⁻□ mg gDW⁻¹, or below 5% of the mean usage) were classed as having a uniquely determined, or forced, commitment; the remainder, whose usage varied across optima, as discretionary.

Roughly 40% of enzyme usages were uniquely determined — a fraction that held steady across the light gradient (Fig. 7). Among enzymes carrying a measured abundance, the forced fraction was 44%, 41%, and 43% at 27.5, 440, and 1100 µmol photons m⁻² s⁻¹ respectively — that is, roughly three-fifths of individual enzyme commitments could take a range of values consistent with the same optimal growth and protein budget. This degeneracy is a normal feature of enzyme-constrained models as many enzymes can substitute for one another. Therefore, individual enzyme or reaction values from a single solution should not be over-interpreted, even though the overall proteome allocation is well constrained.

**Fig. 7.**
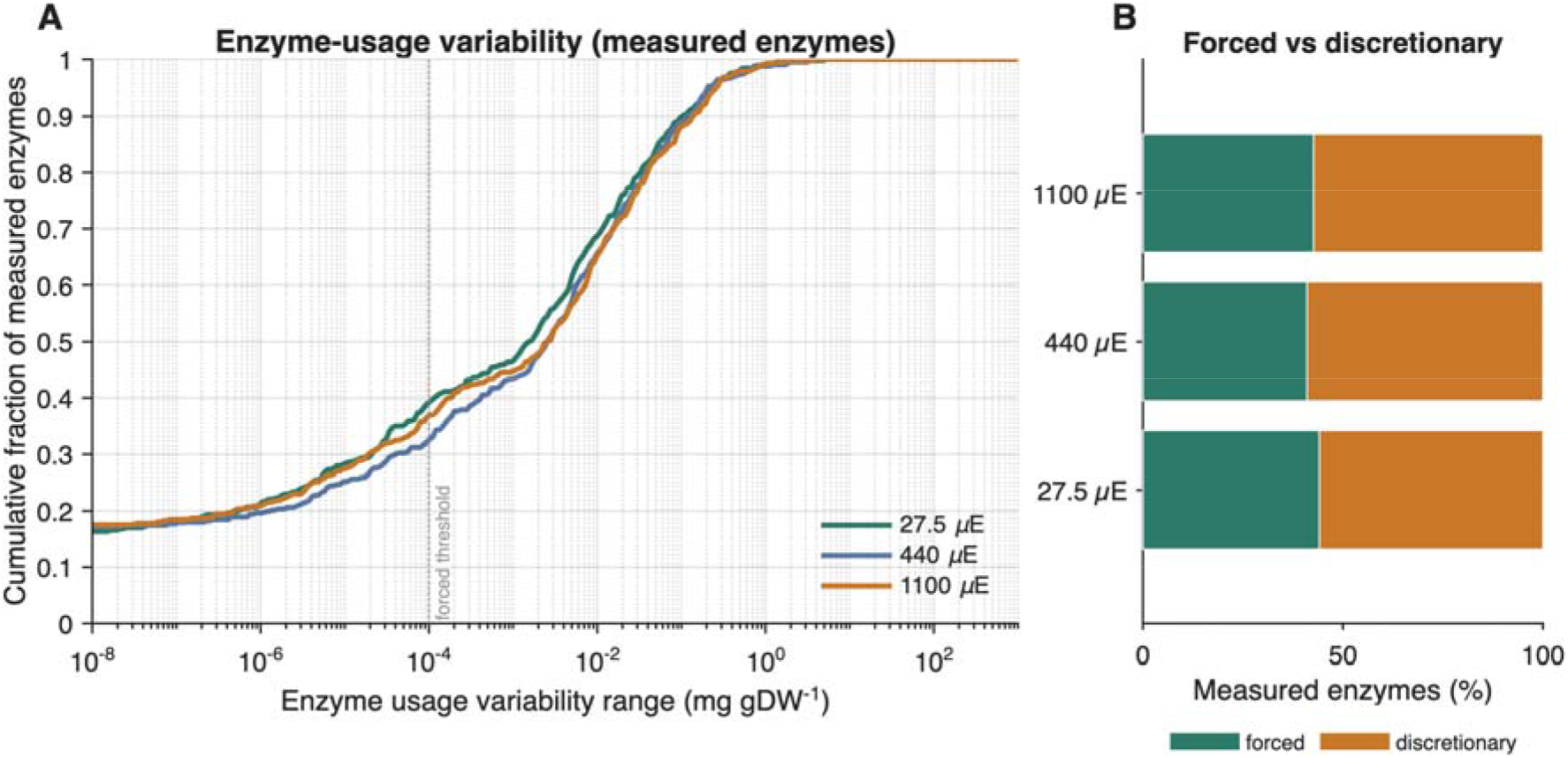
Enzyme-usage variability across the light gradient. Flux variability analysis of the enzyme-usage variables of each condition-specific proteomics-integrated model, with growth fixed at its maximum. For each enzyme the variability range (maximum − minimum usage across alternative optimal solutions) was computed. **(A)** Cumulative distribution of the usage-variability range for enzymes carrying a measured abundance, at 27.5, 440, and 1100 µmol photons m⁻² s⁻¹; the dashed line marks the threshold below which a usage is treated as uniquely determined (“forced”). Curves rising steeply at small ranges indicate that many enzymes are tightly determined, while the long right-hand tails show the enzymes whose usage is free to vary. **(B)** Fraction of measured enzymes with a forced versus discretionary usage, per condition. The forced fraction is stable at ∼41–44% across all three conditions, indicating a structural property of the network rather than a condition-specific effect. Enzymes are classed as forced when their usage range is below 10⁻□ mg gDW⁻¹ or below 5% of the mean usage.

## Discussion

This study analyses the effect of light intensity on the metabolism of *Synechocystis* sp. PCC 6803 using an enzyme-constrained genome-scale model across three acclimated states: light-limited (27.5 µmol photons m⁻² s⁻¹), light-saturated (440 µmol photons m⁻² s⁻¹) and photoinhibited (1100 µmol photons m⁻² s⁻¹). Incorporating enzymatic constraints improved the model’s behaviour over the conventional GEM and, in particular, allowed it to reproduce the decline in growth rate at the highest light intensity, the hallmark of photoinhibition, which stoichiometry alone cannot capture.

In the model this decline arises because growth becomes limited by the finite pool of catalytic protein: once the enzyme budget is fully committed, additional photon supply cannot be converted into faster growth. This resource-limitation interpretation is consistent with quantitative proteomics of *Synechocystis*, which shows that its proteome is a finite, growth-rate-dependent resource and that the maximum growth rate is constrained by proteome allocation [9,30]. Adding enzymatic constraints also reduced the space of feasible flux distributions, by up to two orders of magnitude relative to the conventional GEM, with the reaction population shifting towards the fixed class. Comparable reductions have been reported for the earlier *Synechocystis* ecModel [35] and for enzyme-constrained models of other organisms built within the same framework [29,32–34].

The proteome allocation itself was dominated by the same four subsystems — oxidative phosphorylation, transport, photosynthesis and carbon fixation — at every light intensity, and the total catalytic-protein commitment tracked growth rate rather than light, being largest at the light-saturated condition. This follows from the structure of the model: because enzyme usage is proportional to flux, the protein commitment scales with growth rate. Light itself enters only as the photon-uptake constraint, while the size of the protein pool is set separately from the measured proteome; neither directly imposes a light-dependent proteome structure. Capturing how light shapes the proteome independently of growth lies beyond an enzyme-constrained model of this kind, and calls instead for frameworks in which the light response is represented explicitly, such as coarse-grained proteome-allocation models, which reproduce the light-limited, light-saturated and photoinhibited growth regimes through the cost of the cellular machinery that limits growth [36,37], and, for the underlying redox and photoprotective response, kinetic models in which the alternative electron sinks are treated as free processes rather than fixed constraints [38].

A variability analysis of enzyme usage further showed that fewer than half of individual usages were uniquely determined, the remainder varying freely across equally optimal solutions, so the allocation is well constrained in aggregate but not enzyme by enzyme.

The model represents photoinhibition as a consequence of the finite catalytic-protein budget: at the highest light intensity, the fully committed protein pool, rather than photon supply, limits growth. It does not, however, resolve the biochemical mechanisms of high-light stress. The enzyme constraints in a GECKO-type ecModel apply only to metabolic enzymes; the translational and chaperone machinery and the structural photosynthetic proteins are not represented as individual protein species subject to the enzyme constraints. The turnover of the photosystem II reaction-centre protein D1 — the most rapidly turned-over protein under photodamage, whose degradation by the FtsH protease and resynthesis together consume of the order of 10³ ATP equivalents per molecule [39] — is therefore not resolved: individual structural proteins are not tracked as distinct species with their own synthesis and degradation, and the model has no photodamaged-protein state through which photo-inhibitory turnover could be expressed. Photoinhibition also involves regulated and dynamic processes — the balance between light-induced damage to photosystem II and its repair, and the regulated dissipation of excess reducing equivalents through the photoreduction of O₂ downstream of photosystem I [40]. A steady-state, constraint-based model describes the stoichiometry and enzyme capacity of metabolism rather than its regulation or dynamics and therefore cannot represent these mechanisms. The model thus accounts for the growth-level and resource consequences of photoinhibition, but not the regulatory photo-physiology that produces them.

Several further limitations temper these conclusions. The availability of experimentally measured turnover numbers for *Synechocystis* is limited, and most reported values were obtained in vitro and may not reflect activity in the living cell. Deep-learning prediction of turnover numbers from protein sequence, as integrated in GECKO 3 [4], mitigates the scarcity of measured values, but the in-vivo accuracy of these predictions remains to be validated. The reconstruction and calibration of the model also depend on the quality of the proteomics data: where abundances are missing or imprecise, the model may relax individual enzyme constraints to reach the measured growth rate, and such relaxed or default-bounded enzymes cannot support quantitative claims about their usage, a caution reinforced by the identifiability analysis above. Accurate subunit stoichiometry is likewise required for multi-subunit complexes such as the light-harvesting and photosystem proteins, since the number of subunits scales the calculated usage; the limited coverage of cyanobacterial complexes in resources such as the Complex Portal therefore remains a source of uncertainty.

Future work could address these limitations through fine-grained resource-allocation models, such as Resource Balance Analysis (RBA) [41] Metabolism and Expression (ME) models, [42], and Expression and Thermodynamics Flux (ETFL) models [43], which, unlike enzyme-constrained models, explicitly represent the synthesis and degradation of macromolecules such as enzymes and ribosomes. Kinetic models could further capture the regulation and dynamics that a steady-state description omits. Reconstructing such models for *Synechocystis* would extend the present analysis towards a mechanistic account of how light shapes its metabolism.

## Conclusions

In conclusion, we reconstructed an enzyme-constrained genome-scale model of *Synechocystis* sp. PCC 6803 and used it to study metabolism and proteome allocation across three light intensities. By incorporating enzymatic constraints, the model reproduced the decline in growth under high light, which the conventional genome-scale model cannot capture, attributing it to the depletion of the finite catalytic-protein. Integrating absolute proteomics further constrained the model: enzyme constraints progressively reduced the feasible flux space, and the proteome allocation was dominated by a small number of subsystems that tracked growth rate across conditions. At the same time, flux variability analysis showed that fewer than half of individual enzyme usages took a single, well-defined value, the rest varying across alternative optimal solutions. The model captures the growth and resource consequences of growth at different light intensities, but not the regulatory and dynamic physiology underlying the high-light response, the latter would require fine-grained resource-allocation models (RBA, ME) to represent protein turnover and the biosynthetic machinery, and kinetic models to represent regulation and dynamics. Together, these results contribute to enzyme-constrained modelling of cyanobacterial metabolism and its use in understanding responses to light.

## Code Availability

The source code and analysis scripts supporting this work are currently maintained in a private repository and will be made publicly available upon submission to a peer-reviewed journal.

