## Supplemental Data 1 for "Light-dependent metabolism and proteome allocation in *Synechocystis* sp. PCC 6803: an enzyme-constrained metabolic model across light intensities"

### Supplementary Information

**Table S1.** Light-condition-specific macromolecular composition of *Synechocystis* sp. PCC 6803 used to construct the biomass reactions (Zavřel et al. [1]).

| Light (μmol photons $m^{-2}$ $s^{-1}$) | μ ($h^{-1}$) | Protein (mg $gDW^{-1}$) | Glycogen (mg $gDW^{-1}$) | Chlorophyll a (mg $gDW^{-1}$) | Carotenoids (mg $gDW^{-1}$) |
|---|---|---|---|---|---|
| 27.5 | 0.0254 | 402.5 | 83.8 | 16.0 | 4.4 |
| 440 | 0.1044 | 278.9 | 192.2 | 7.8 | 2.8 |
| 1100 | 0.0933 | 264.0 | 228.6 | 5.8 | 2.9 |

**Table S2.** Curated turnover numbers (Kcat) for the *Synechocystis* sp. PCC 6803 ecModel, with evidence source and fold-change per reaction

| Reaction | Kcat, old ($s^{-1}$) | Kcat, new ($s^{-1}$) | Source | Fold change | UniProt ID | Reaction formula |
|---|---|---|---|---|---|---|
| CHORS | 0.0083 | 16.1 | DLKcat | 1940 | P23353 | 5-O-(1-Carboxyvinyl)-3-phosphoshikimate[c] + 2.706 prot_P23353[c] => Chorismate[c] + Phosphate[c] |
| 3OAS100_EXP_4 | 0.047 | 8.57 | KcatNet | 182 | P74017 | H+[c] + Malonyl-[acyl-carrier protein][c] + Octanoyl-ACP (n-C8:0ACP)[c] + 1.4011 prot_P74017[c] => 3-Oxodecanoyl-[acyl-carrier protein][c] + Acyl carrier protein[c] + CO2 CO2[c] |
| 3OAS60_EXP_2 | 0.047 | 8.57 | KcatNet | 182 | P74017 | Butyryl-ACP (n-C4:0ACP)[c] + H+[c] + Malonyl-[acyl-carrier protein][c] + 1.4011 prot_P74017[c] => 3-Oxohexanoyl-[acyl-carrier protein][c] + Acyl carrier protein[c] + CO2 CO2[c] |
| 3OAS80_EXP_2 | 0.047 | 8.57 | KcatNet | 182 | P74017 | H+[c] + Hexanoyl-ACP (n-C6:0ACP)[c] + Malonyl-[acyl-carrier protein][c] + 1.4011 prot_P74017[c] => 3-Oxooctanoyl-[acyl-carrier protein][c] + Acyl carrier protein[c] + CO2 CO2[c] |
| 3OAS120_EXP_4 | 0.047 | 8.57 | KcatNet | 182 | P74017 | Decanoyl-ACP (n-C10:0ACP)[c] + H+[c] + Malonyl-[acyl-carrier protein][c] + 1.4011 prot_P74017[c] => 3-Oxododecanoyl-[acyl-carrier protein][c] + Acyl carrier protein[c] + CO2 CO2[c] |
| 3OAS140_EXP_4 | 0.047 | 8.57 | KcatNet | 182 | P74017 | Dodecanoyl-ACP (n-C12:0ACP)[c] + H+[c] + Malonyl-[acyl-carrier protein][c] + 1.4011 prot_P74017[c] => 3-Oxotetradecanoyl-[acyl-carrier protein][c] + Acyl carrier protein[c] + CO2 CO2[c] |
| 3OAS160_EXP_4 | 0.047 | 8.57 | KcatNet | 182 | P74017 | H+[c] + Malonyl-[acyl-carrier protein][c] + Myristoyl-ACP (n-C14:0ACP)[c] + 1.4011 prot_P74017[c] => 3-Oxohexadecanoyl-[acyl-carrier protein][c] + Acyl carrier protein[c] + CO2 CO2[c] |

| Reaction | Kcat, old ($s^{-1}$) | Kcat, new ($s^{-1}$) | Source | Fold change | UniProt ID | Reaction formula |
|---|---|---|---|---|---|---|
| FCLT | 0.0002 | 0.613 | KcatNet | 3070 | P54225 | Fe2+ mitochondria[c] + Protoporphyrin[c] + 19.8893 prot_P54225[c] => 2 H+[c] + Protoheme C34H30FeN4O4[c] |
| 3OAS180_EXP_2 | 0.047 | 8.57 | KcatNet | 182 | P74017 | H+[c] + Malonyl-[acyl-carrier protein][c] + Palmitoyl-ACP (n-C16:0ACP)[c] + 1.4011 prot_P74017[c] => 3-Oxooctadecanoyl-[acyl-carrier protein][c] + Acyl carrier protein[c] + CO2 CO2[c] |
| TMDS | 0.0072 | 1.06 | KcatNet | 147 | P73053 | DUMP C9H11N2O8P[c] + 5,10-Methylenetetrahydrofolate[c] + 26.2478 prot_P73053[c] => 7,8-Dihydrofolate[c] + DTMP C10H13N2O8P[c] |
| DES_6B | 0.0378 | 4.55 | KcatNet | 120 | Q08871 | 2 Reduced ferredoxin[c] + 2 H+[c] + O2 O2[c] + Linoleoyl-ACP (n-C18 2ACP)[c] + 2.5292 prot_Q08871[c] => 2 Oxidized ferredoxin[c] + 2 H2O H2O[c] + G-linolenoilACP[c] |
| 3OAS121_EXP_2 | 0.047 | 8.57 | KcatNet | 182 | P74017 | Cis-dec-3-enoyl-[acyl-carrier protein] (n-C10:1)[c] + H+[c] + Malonyl-[acyl-carrier protein][c] + 1.4011 prot_P74017[c] => 3-oxo-cis-dodec-5-enoyl-[acyl-carrier protein][c] + Acyl carrier protein[c] + CO2 CO2[c] |
| 3OAS141_EXP_2 | 0.047 | 8.57 | KcatNet | 182 | P74017 | Cis-dodec-5-enoyl-[acyl-carrier protein] (n-C12:1)[c] + H+[c] + Malonyl-[acyl-carrier protein][c] + 1.4011 prot_P74017[c] => 3-oxo-cis-myristol-7-eoyl-[acyl-carrier protein][c] + Acyl carrier protein[c] + CO2 CO2[c] |
| 3OAS161_EXP_4 | 0.047 | 8.57 | KcatNet | 182 | P74017 | H+[c] + Malonyl-[acyl-carrier protein][c] + Cis-tetradec-7-enoyl-[acyl-carrier protein] (n-C14:1)[c] + 1.4011 prot_P74017[c] => 3-oxo-cis-palm-9-eoyl-[acyl-carrier protein][c] + Acyl carrier protein[c] + CO2 CO2[c] |
| CPPPGO | 0.017 | 0.541 | DLKcat | 31.8 | P72848 | Coproporphyrinogen III[c] + 2 H+[c] + O2 O2[c] + 39.992 prot_P72848[c] => 2 CO2 CO2[c] + 2 H2O H2O[c] + Protoporphyrinogen IX[c] |
| MPML | 0.0188 | 4.74 | DLKcat | 252 | P51634 | ATP C10H12N5O13P3[c] + H2O H2O[c] + Magnesium[c] + Protoporphyrin[c] + 2.3141 prot_P51634[c] + 4.3196 prot_P72772[c] + 8.7111 prot_P73020[c] => ADP C10H12N5O10P2[c] + 3 H+[c] + Magnesium protoporphyrin[c] + Phosphate[c] |
| PAPSR | 0.335 | 5.55 | KcatNet | 16.6 | P72794 | 3'-Phosphoadenylyl sulfate[c] + Reduced thioredoxin[c] + 1.4256 prot_P72794[c] => 2 H+[c] + Adenosine 3',5'-bisphosphate[c] + Sulfite[c] + Oxidized thioredoxin[c] |
| 3OAS181_EXP_4 | 0.047 | 8.57 | KcatNet | 182 | P74017 | H+[c] + Cis-hexadec-9-enoyl-[acyl-carrier protein] (n-C16:1)[c] + Malonyl-[acyl-carrier protein][c] + 1.4011 prot_P74017[c] => 3-oxo- |

| Reaction | Kcat, old ($s^{-1}$) | Kcat, new ($s^{-1}$) | Source | Fold change | UniProt ID | Reaction formula |
|---|---|---|---|---|---|---|
| | | | | | | cis-vacc-11-enoyl-[acyl-carrier protein][c] + Acyl carrier protein[c] + CO2 CO2[c] |
| ARGSS | 0.687 | 8.28 | KcatNet | 12.1 | P77973 | L-Aspartate[c] + ATP C10H12N5O13P3[c] + L-Citrulline[c] + 5.9677 prot_P77973[c] => AMP C10H12N5O7P[c] + N(omega)-(L-Arginino)succinate[c] + H+[c] + Diphosphate[c] |
| PGMT_REV_EXP_1 | 11.3 | 36 | KcatNet | 3.18 | P74643 | D-Glucose 6-phosphate[c] + 0.47116 prot_P74643[c] => D-Glucose 1-phosphate[c] |
| CDPMEK | 0.75 | 3.89 | DLKcat | 5.19 | P72663 | 4-(cytidine 5'-diphospho)-2-C-methyl-D-erythritol[c] + ATP C10H12N5O13P3[c] + 2.4321 prot_P72663[c] => 2-phospho-4-(cytidine 5'-diphospho)-2-C-methyl-D-erythritol[c] + ADP C10H12N5O10P2[c] + H+[c] |
| UGMDDS | 0.355 | 3.02 | KcatNet | 8.49 | P45450 | D-Alanyl-D-alanine[c] + ATP C10H12N5O13P3[c] + UDP-N-acetylmuramoyl-L-alanyl-D-gamma-glutamyl-meso-2,6-diaminopimelate[c] + 4.4448 prot_P45450[c] => ADP C10H12N5O10P2[c] + H+[c] + Phosphate[c] + UDP-N-acetylmuramoyl-L-alanyl-D-glutamyl-meso-2,6-diaminopimeloyl-D-alanyl-D-alanine[c] |
| BPNT | 1.12 | 4.89 | KcatNet | 4.36 | Q55507 | H2O H2O[c] + Adenosine 3',5'-bisphosphate[c] + 1.8275 prot_Q55507[c] => AMP C10H12N5O7P[c] + Phosphate[c] |
| G3PAT160_EXP_2 | 0.831 | 4.8 | KcatNet | 5.78 | P73933 | Glycerol 3-phosphate[c] + Palmitoyl-ACP (n-C16:0ACP)[c] + 1.3609 prot_P73933[c] => 1-hexadecanoyl-sn-glycerol 3-phosphate[c] + Acyl carrier protein[c] |
| DES_6 | 0.0378 | 4.55 | KcatNet | 120 | Q08871 | 2 Reduced ferredoxin[c] + 2 H+[c] + O2 O2[c] + A-lineloyl-ACP (n-C18 3ACP)[c] + 2.5292 prot_Q08871[c] => 2 Oxidized ferredoxin[c] + 2 H2O H2O[c] + OctadecatetraenoilACP[c] |
| SHCHCS2 | 0.05 | 0.747 | KcatNet | 14.9 | Q55725 | Isochorismate[c] + Succinate semialdehyde-thiamin diphosphate anion[c] + 49.156 prot_Q55725[c] => 2-Succinyl-6-hydroxy-2,4-cyclohexadiene-1-carboxylate[c] + Pyruvate[c] + Thiamine diphosphate[c] |
| PRAGSr | 7 | 21.1 | KcatNet | 3.01 | P74232 | ATP C10H12N5O13P3[c] + Glycine[c] + 5-Phospho-beta-D-ribosylamine[c] + 0.5802 prot_P74232[c] => ADP C10H12N5O10P2[c] + N1-(5-Phospho-D-ribosyl)glycinamide[c] + H+[c] + Phosphate[c] |
| GLUTRS | 2.15 | 9.72 | KcatNet | 4.52 | Q55778 | ATP C10H12N5O13P3[c] + L-Glutamate[c] + TRNA (Glu)[c] + 1.5429 prot_Q55778[c] => AMP C10H12N5O7P[c] + L-Glutamyl-tRNA(Glu)[c] + Diphosphate[c] |

| Reaction | Kcat, old ($s^{-1}$) | Kcat, new ($s^{-1}$) | Source | Fold change | UniProt ID | Reaction formula |
|---|---|---|---|---|---|---|
| POR_1_EXP_2 | 0.165 | 4.32 | DLKcat | 26.2 | Q59987 | H+[c] + Nicotinamide adenine dinucleotide phosphate - reduced[c] + Protochlorophyllide[c] + 2.3199 prot_Q59987[c] => Chlorophyllide a[c] + Nicotinamide adenine dinucleotide phosphate[c] |
| PGSA160 | 0.102 | 1.45 | KcatNet | 14.2 | P74372 | CDP-1,2-dihexadecanoylglycerol[c] + Glycerol 3-phosphate[c] + 3.7782 prot_P74372[c] => CMP C9H12N3O8P[c] + H+[c] + Phosphatidylglycerophosphate (dihexadecanoyl, n-C16:0)[c] |
| GHMT2r_REV | 5 | 16.6 | DLKcat | 3.32 | P77962 | Glycine[c] + H2O H2O[c] + 5,10-Methylenetetrahydrofolate[c] + 1.5477 prot_P77962[c] => L-Serine[c] + 5,6,7,8-Tetrahydrofolate[c] |
| CHPHYS | 0.186 | 2.18 | KcatNet | 11.7 | Q55145 | Chlorophyllide a[c] + H+[c] + Phytyl diphosphate[c] + 4.4836 prot_Q55145[c] => Chlorophyll a[c] + Diphosphate[c] |
| UAMAS | 1 | 3.48 | DLKcat | 3.48 | P74528 | L-Alanine[c] + ATP C10H12N5O13P3[c] + UDP-N-acetylmuramate[c] + 4.3559 prot_P74528[c] => ADP C10H12N5O10P2[c] + H+[c] + Phosphate[c] + UDP-N-acetylmuramoyl-L-alanine[c] |
| GLUTRR | 2.52 | 13.5 | KcatNet | 5.38 | P28463 | L-Glutamyl-tRNA(Glu)[c] + H+[c] + Nicotinamide adenine dinucleotide phosphate - reduced[c] + 1.9515 prot_P28463[c] => L-Glutamate 1-semialdehyde[c] + Nicotinamide adenine dinucleotide phosphate[c] + TRNA (Glu)[c] |
| HMBS | 0.25 | 31.8 | KcatNet | 127 | P73660 | H2O H2O[c] + 4 Porphobilinogen[c] + 0.30479 prot_P73660[c] => Hydroxymethylbilane[c] + 4 Ammonium[c] |
| UPPDC1 | 0.317 | 3.33 | KcatNet | 10.5 | P54224 | 4 H+[c] + Uroporphyrinogen III[c] + 6.5361 prot_P54224[c] => 4 CO2 CO2[c] + Coproporphyrinogen III[c] |
| PGSA182_9_12 | 0.102 | 1.45 | KcatNet | 14.2 | P74372 | CDP-1,2-dioctadec-9-12-dienoylglycerol[c] + Glycerol 3-phosphate[c] + 3.7782 prot_P74372[c] => CMP C9H12N3O8P[c] + H+[c] + Phosphatidylglycerophosphate (dioctadec-9-12-dienoyl, n-C18 2)[c] |
| AGPR_REV | 5.34 | 20 | KcatNet | 3.74 | P54899 | N-Acetyl-L-glutamyl 5-phosphate[c] + H+[c] + Nicotinamide adenine dinucleotide phosphate - reduced[c] + 0.531 prot_P54899[c] => N-Acetyl-L-glutamate 5-semialdehyde[c] + Nicotinamide adenine dinucleotide phosphate[c] + Phosphate[c] |
| MTAP | 0.26 | 2.18 | DLKcat | 8.39 | P0DJF8 | 5-Methylthioadenosine[c] + Phosphate[c] + 26.9862 prot_P0DJF8[c] => 5-Methylthio-5-deoxy-D-ribose 1-phosphate[c] + Adenine[c] |
| NPHS | 0.062 | 7.03 | KcatNet | 113 | P73495 | O-Succinylbenzoyl-CoA[c] + 1.1971 prot_P73495[c] => Coenzyme A[c] + 1,4-Dihydroxy-2-naphthoate[c] |

| Reaction | Kcat, old ($s^{-1}$) | Kcat, new ($s^{-1}$) | Source | Fold change | UniProt ID | Reaction formula |
|---|---|---|---|---|---|---|
| G3PAT182_9_12_EXP_2 | 0.831 | 4.8 | KcatNet | 5.78 | P73933 | Glycerol 3-phosphate[c] + Linoleoyl-ACP (n-C18 2ACP)[c] + 1.3609 prot_P73933[c] => 1-octadec-9-12-dienoyl-sn-glycerol 3-phosphate[c] + Acyl carrier protein[c] |
| G3PAT184_6_9_12_15_EXP_2 | 0.831 | 4.8 | KcatNet | 5.78 | P73933 | Glycerol 3-phosphate[c] + G-linolenoilACP[c] + 1.3609 prot_P73933[c] => 1-octadec-6-9-12trienoyl-sn-glycerol 3-phosphate[c] + Acyl carrier protein[c] |
| FRTT | 1.1 | 5.16 | DLKcat | 4.69 | P72683 | Farnesyl diphosphate[c] + Isopentenyl diphosphate[c] + 1.7426 prot_P72683[c] => Geranylgeranyl diphosphate C20H33O7P2[c] + Diphosphate[c] |
| HGPHT | 0.0352 | 7.55 | KcatNet | 214 | P73726 | H+[c] + Homogentisate C8H7O4[c] + Phytyl diphosphate[c] + 1.266 prot_P73726[c] => 2-Methyl-6-phytylquinol[c] + CO2 CO2[c] + Diphosphate[c] |
| GRTT | 1.5 | 15 | DLKcat | 10.1 | P37294 | Geranyl diphosphate[c] + Isopentenyl diphosphate[c] + 0.71465 prot_P37294[c] => Farnesyl diphosphate[c] + Diphosphate[c] |
| PPND | 3.53 | 12.3 | KcatNet | 3.48 | P73906 | Nicotinamide adenine dinucleotide[c] + Prephenate[c] + 0.6815 prot_P73906[c] => 3-(4-Hydroxyphenyl)pyruvate[c] + CO2 CO2[c] + Nicotinamide adenine dinucleotide - reduced[c] |
| UHGADA | 3.3 | 18.5 | KcatNet | 5.61 | P72988 | H2O H2O[c] + UDP-3-O-(3-hydroxytetradecanoyl)-N-acetylglucosamine[c] + 0.45515 prot_P72988[c] => Acetate[c] + UDP-3-O-(3-hydroxytetradecanoyl)-D-glucosamine[c] |
| PHYTEDH2 | 0.546 | 3 | KcatNet | 5.49 | P29273 | Nicotinamide adenine dinucleotide phosphate[c] + All-trans-Phytoene[c] + 4.9041 prot_P29273[c] => H+[c] + Nicotinamide adenine dinucleotide phosphate - reduced[c] + All-trans-Phytofluene[c] |
| PHYTFDH2 | 0.546 | 3 | KcatNet | 5.49 | P29273 | Nicotinamide adenine dinucleotide phosphate[c] + All-trans-Phytofluene[c] + 4.9041 prot_P29273[c] => H+[c] + Nicotinamide adenine dinucleotide phosphate - reduced[c] + Zeta-Carotene[c] |
| RBFSa | 0.04 | 31.2 | KcatNet | 781 | P73527 | 4-(1-D-Ribitylamino)-5-aminouracil[c] + 3,4-dihydroxy-2-butanone 4-phosphate[c] + 0.15672 prot_P73527[c] => 6,7-Dimethyl-8-(1-D-ribityl)lumazine[c] + 2 H2O H2O[c] + Phosphate[c] |
| OXGDC2 | 0.28 | 1.12 | KcatNet | 4 | Q55725 | 2-Oxoglutarate[c] + H+[c] + Thiamine diphosphate[c] + 32.8076 prot_Q55725[c] => CO2 CO2[c] + Succinate semialdehyde-thiamin diphosphate anion[c] |
| SUCBZL | 0.317 | 1.84 | KcatNet | 5.82 | Q55182 | ATP C10H12N5O13P3[c] + Coenzyme A[c] + O-Succinylbenzoate[c] + 8.3184 prot_Q55182[c] => AMP C10H12N5O7P[c] + Diphosphate[c] + O-Succinylbenzoyl-CoA[c] |

| Reaction | Kcat, old ($s^{-1}$) | Kcat, new ($s^{-1}$) | Source | Fold change | UniProt ID | Reaction formula |
|---|---|---|---|---|---|---|
| UAMAGS | 4.33 | 18.9 | KcatNet | 4.37 | P73668 | ATP C10H12N5O13P3[c] + D-Glutamate[c] + UDP-N-acetylmuramoyl-L-alanine[c] + 0.71953 prot_P73668[c] => ADP C10H12N5O10P2[c] + H+[c] + Phosphate[c] + UDP-N-acetylmuramoyl-L-alanyl-D-glutamate[c] |
| PGSA183_9_12_15 | 0.102 | 1.45 | KcatNet | 14.2 | P74372 | CDP-1,2-dioctadec-9-12-15-trienoylglycerol[c] + Glycerol 3-phosphate[c] + 3.7782 prot_P74372[c] => CMP C9H12N3O8P[c] + H+[c] + Phosphatidylglycerophosphate (dioctadec-9-12-15-trienoyl, n-C18 3)[c] |
| MTRI | 1.07 | 3.87 | KcatNet | 3.61 | P74497 | 5-Methylthio-5-deoxy-D-ribose 1-phosphate[c] + 2.748 prot_P74497[c] => 5-Methylthio-5-deoxy-D-ribulose 1-phosphate[c] |
| DHFS | 0.124 | 3.72 | KcatNet | 29.9 | P73248 | ATP C10H12N5O13P3[c] + Dihydropteroate[c] + L-Glutamate[c] + 2.359 prot_P73248[c] => ADP C10H12N5O10P2[c] + 7,8-Dihydrofolate[c] + H+[c] + Phosphate[c] |
| G3PAT161_EXP_2 | 0.831 | 4.8 | KcatNet | 5.78 | P73933 | Glycerol 3-phosphate[c] + Cis-hexadec-9-enoyl-[acyl-carrier protein] (n-C16:1)[c] + 1.3609 prot_P73933[c] => 1-hexadec-9-enoyl-sn-glycerol 3-phosphate[c] + Acyl carrier protein[c] |
| UAAGDS | 8.77 | 30.1 | DLKcat | 3.44 | Q55469 | Meso-2,6-Diaminoheptanedioate[c] + ATP C10H12N5O13P3[c] + UDP-N-acetylmuramoyl-L-alanyl-D-glutamate[c] + 0.50346 prot_Q55469[c] => ADP C10H12N5O10P2[c] + H+[c] + Phosphate[c] + UDP-N-acetylmuramoyl-L-alanyl-D-gamma-glutamyl-meso-2,6-diaminopimelate[c] |
| PC20M | 0.0401 | 6.66 | KcatNet | 166 | P73644 | S-Adenosyl-L-methionine[c] + Dihydrosirohydrochlorin[c] + 1.1257 prot_P73644[c] => S-Adenosyl-L-homocysteine[c] + H+[c] + Precorrin 3 A[c] |
| G3PAT181_9_EXP_2 | 0.831 | 4.8 | KcatNet | 5.78 | P73933 | Glycerol 3-phosphate[c] + Cis-octadec-9-enoyl-[acyl-carrier protein] (n-C18 1)[c] + 1.3609 prot_P73933[c] => 1-octadec-9-enoyl-sn-glycerol 3-phosphate[c] + Acyl carrier protein[c] |
| G3PAT181_EXP_2 | 0.831 | 4.8 | KcatNet | 5.78 | P73933 | Glycerol 3-phosphate[c] + Cis-octadec-11-enoyl-[acyl-carrier protein] (n-C18:1)[c] + 1.3609 prot_P73933[c] => 1-octadec-11-enoyl-sn-glycerol 3-phosphate[c] + Acyl carrier protein[c] |
| DHNPA_1 | 0.089 | 7.06 | KcatNet | 79.3 | P74342 | Dihydroneopterin[c] + 0.5171 prot_P74342[c] => 2 Amino 4 hydroxy 6 hydroxymethyl 7 8 dihydropteridine C7H9N5O2[c] + Glycolaldehyde[c] |
| PHYTES2 | 1.42 | 5.27 | DLKcat | 3.7 | P37294 | Prephytoene diphosphate[c] + 2.0401 prot_P37294[c] => All-trans-Phytoene[c] + Diphosphate[c] |

| Reaction | Kcat, old ($s^{-1}$) | Kcat, new ($s^{-1}$) | Source | Fold change | UniProt ID | Reaction formula |
| --- | --- | --- | --- | --- | --- | --- |
| QULNS | 0.8 | 3.81 | DLKcat | 4.76 | P74578 | Dihydroxyacetone phosphate[c] + Iminoaspartate[c] + 2.5834 prot_P74578[c] => 2 H2O H2O[c] + Phosphate[c] + Quinolinate[c] |
| PGSA181_9 | 0.102 | 1.45 | KcatNet | 14.2 | P74372 | CDP-1,2-dioctadec-9-enoylglycerol [c] + Glycerol 3-phosphate[c] + 3.7782 prot_P74372[c] => CMP C9H12N3O8P[c] + H+[c] + Phosphatidylglycerophosphate (dioctadec-9-enoyl, n-C18 1)[c] |
| THRPDC | 0.105 | 1.06 | KcatNet | 10.2 | P73417 | H+[c] + L-Threonine O-3-phosphate[c] + 10.1601 prot_P73417[c] => D-1-Aminopropan-2-ol O-phosphate[c] + CO2 CO2[c] |
| H2CO3_NAt_syn_EXP_1 | 2.21 | 15.1 | KcatNet | 6.84 | P73953 | Bicarbonate[p] + Sodium[p] + 0.7291 prot_P73953[c] => Bicarbonate[c] + Sodium[c] |
| HEMEOS | 0.203 | 2.49 | KcatNet | 12.3 | Q79EF2 | Farnesyl diphosphate[c] + H2O H2O[c] + Protoheme C34H30FeN4O4[c] + 3.8915 prot_Q79EF2[c] => Heme O C49H56FeN4O5[c] + Diphosphate[c] |
| R05224_1 | 0.16 | 5.6 | KcatNet | 35 | P73002 | 2 ATP C10H12N5O13P3[c] + Reduced ferredoxin[c] + 2 L-Glutamine[c] + H2O H2O[c] + Hydrogenobyrinate[c] + 2.669 prot_P73002[c] => 2 ADP C10H12N5O10P2[c] + Oxidized ferredoxin[c] + 2 L-Glutamate[c] + H+[c] + Hydrogenobyrinate a,c diamide[c] + Diphosphate[c] |
| FBA_REV_EXP_1 | 1.49 | 14.9 | sensitivityTuning | 10 | P74309 | Dihydroxyacetone phosphate[c] + Glyceraldehyde 3-phosphate[c] + 0.61874 prot_P74309[c] => D-Fructose 1,6-bisphosphate[c] |
| FBP_EXP_2 | 10.5 | 105 | sensitivityTuning | 10 | P73922 | D-Fructose 1,6-bisphosphate[c] + H2O H2O[c] + 0.39233 prot_P73922[c] => D-Fructose 6-phosphate[c] + Phosphate[c] |
| H2Otu_syn_REV | 20.9 | 209 | sensitivityTuning | 10 | P73809 | H2O H2O[c] + 0.13582 prot_P73809[c] => H2O H2O[u] |
| PSII_EXP_4 | 668 | 6680 | sensitivityTuning | 10 | Q55013 | H2O H2O[u] + 2 H+[c] + 2 Light[c] + Plastoquinone[u] + 0.0046494 prot_P05429[c] + 0.0032977 prot_P07826[c] + 0.00078586 prot_P09190[c] + 0.00041027 prot_P09191[c] + 0.0032846 prot_P09192[c] + 0.0041836 prot_P09193[c] + 0.0024877 prot_P10549[c] + 0.00059183 prot_P14835[c] + 0.00042516 prot_P15819[c] + 0.00018763 prot_P26286[c] + 0.00034814 prot_P72575[c] + 0.00096388 prot_P72652[c] + 0.00032286 prot_P72701[c] + 0.0067079 prot_P72870[c] + 0.00032835 prot_P73070[c] + 0.0072112 prot_P73093[c] + 0.0023259 prot_P73202[c] + 0.0081142 prot_P73203[c] + 0.007684 prot_P73204[c] + 0.0012223 prot_P73638[c] + 0.00034948 prot_P73676[c] + 0.0011738 prot_P73731[c] + 0.0017255 prot_P73952[c] + 0.0012297 prot_P74367[c] + 0.0023568 prot_P74551[c] + 0.0019227 prot_Q01950[c] + 0.017378 prot_Q01951[c] + 0.023625 prot_Q01952[c] + 0.00035812 |

| Reaction | Kcat, old ($s^{-1}$) | Kcat, new ($s^{-1}$) | Source | Fold change | UniProt ID | Reaction formula |
|---|---|---|---|---|---|---|
| | | | | | | prot_Q54697[c] + 0.081406 prot_Q54714[c] + 0.078985 prot_Q54715[c] + 0.0014874 prot_Q55013[c] + 0.0011847 prot_Q55332[c] + 0.00037201 prot_Q55354[c] + 0.0010471 prot_Q55356[c] + 0.012512 prot_Q55544[c] => 2 H+[u] + 0.5 O2 O2[u] + Plastoquinol[u] |
| PSI_EXP_2 | 387 | 3870 | sensitivityTuning | 10 | P29256 | 2 Oxidized ferredoxin[c] + 2 Plastocyanin(Cu+)[u] + 2 Light[c] + 0.0017539 prot_P12975[c] + 0.0033686 prot_P19569[c] + 0.017862 prot_P29254[c] + 0.017505 prot_P29255[c] + 0.0039296 prot_P29256[c] + 0.0019009 prot_P32422[c] + 0.0035797 prot_P37277[c] + 0.0016637 prot_P72652[c] + 0.0018613 prot_P72712[c] + 0.011578 prot_P72870[c] + 0.00072782 prot_P72986[c] + 0.0040146 prot_P73202[c] + 0.014006 prot_P73203[c] + 0.013263 prot_P73204[c] + 0.0021097 prot_P73638[c] + 0.004068 prot_P74551[c] + 0.012284 prot_P74625[c] + 0.0033187 prot_Q01950[c] + 0.029995 prot_Q01951[c] + 0.040779 prot_Q01952[c] + 0.14051 prot_Q54714[c] + 0.13633 prot_Q54715[c] + 0.00097588 prot_Q55329[c] + 0.00095026 prot_Q55330[c] + 0.021597 prot_Q55544[c] => 2 Reduced ferredoxin[c] + 2 Plastocyanin(Cu2+)[u] |
| RBPC | 14.3 | 143 | sensitivityTuning | 10 | P54205 | CO2 CO2[c] + H2O H2O[c] + D-Ribulose 1,5-bisphosphate[c] + 0.81685 prot_P54205[c] + 0.20602 prot_P54206[c] => 2 3-Phospho-D-glycerate[c] + 2 H+[c] |

**Table S3.** Per-replicate proteomics scaling. For each light condition and biological replicate, the summed protein mass detected by proteomics, the measured total protein content (Ptot, Zavřel et al. [1]), and the scale factor (Ptot / summed proteomics) applied to place abundances on an absolute mass basis.

| Light (µmol photons $m^{-2}$ $s^{-1}$) | Replicate | Summed proteomics (mg $gDW^{-1}$) | Ptot (mg $gDW^{-1}$) | Scale factor |
|---|---|---|---|---|
| 27.5 | A | 164.97 | 325.30 | 1.97 |
| 27.5 | B | 139.05 | 380.30 | 2.74 |
| 27.5 | C | 154.30 | 373.20 | 2.42 |
| 27.5 | D | 142.63 | 399.60 | 2.80 |
| 27.5 | E | 142.26 | 534.10 | 3.75 |
| 440 | A | 98.25 | 199.80 | 2.03 |
| 440 | B | 101.47 | 276.50 | 2.73 |

| Light (μmol photons $m^{-2}$ $s^{-1}$) | Replicate | Summed proteomics (mg $gDW^{-1}$) | Ptot (mg $gDW^{-1}$) | Scale factor |
|---|---|---|---|---|
| 440 | C | 82.63 | 283.30 | 3.43 |
| 440 | D | 85.49 | 326.40 | 3.82 |
| 440 | E | 99.23 | 308.40 | 3.11 |
| 1100 | A | 74.14 | 203.40 | 2.74 |
| 1100 | B | 70.61 | 204.80 | 2.90 |
| 1100 | C | 73.27 | 269.00 | 3.67 |
| 1100 | D | 67.11 | 322.90 | 4.81 |
| 1100 | E | 95.93 | 320.10 | 3.34 |

**Table S4.** Total protein content and catalytic protein fraction (mean ± SD, n = 5).

| Light (μmol photons $m^{-2}$ $s^{-1}$) | Mean Ptot (mg $gDW^{-1}$) | f |
|---|---|---|
| 27.5 | 402.5 | 0.727 ± 0.023 |
| 440 | 278.9 | 0.669 ± 0.046 |
| 1100 | 264.0 | 0.609 ± 0.061 |

**Table S5.** Metabolic constraints applied to the ecModel per light condition. Exchange-flux bounds for photon, bicarbonate, and glucose uptake, and the biomass upper bound (set to the measured growth rate). Uptake fluxes are negative.

| Light condition (μmol photons $m^{-2}$ $s^{-1}$) | Growth rate ($h^{-1}$) | Photon uptake (mmol $gDW^{-1}$ $h^{-1}$) | Bicarbonate uptake (mmol $gDW^{-1}$ $h^{-1}$) | Glucose uptake (mmol $gDW^{-1}$ $h^{-1}$) |
|---|---|---|---|---|
| 27.5 | 0.0254 | −22.56 | −1000 (unbounded) | 0 |
| 440 | 0.1044 | −183.57 | −1000 (unbounded) | 0 |
| 1100 | 0.0933 | −414.37 | −1000 (unbounded) | 0 |

**Table S6.** Photobioreactor light-capture measurements used to derive photon uptake rates for the three modelled light conditions (T. Zavřel, personal communication; not reported in Zavřel et al. [1]). For each condition, the table gives the cuvette geometry, cellular dry weight, the fraction of available light captured by the cells, and the resulting specific photon uptake rate. The final column gives the photon uptake rates from which the model constraints were taken.

| Incident light (µmol photons $m^{-2}$ $s^{-1}$) | Cuvette area ($m^2$) | Cuvette volume (L) | Dry weight (mg $L^{-1}$) | Dry weight (g $cuvette^{-1}$) | Light captured (%) | Captured intensity (µmol photons $m^{-2}$ $s^{-1}$) | Captured (µmol photons $cuvette^{-1}$ $s^{-1}$) | Specific uptake (µmol photons $gDW^{-1}$ $s^{-1}$) | Specific uptake (mmol photons $gDW^{-1}$ $h^{-1}$) |
|---|---|---|---|---|---|---|---|---|---|
| 27.5 | 0.0189 | 0.42 | 132.2 | 0.056 | 67 | 18 | 0.35 | 6 | 22.56 |
| 440 | 0.0189 | 0.42 | 221.4 | 0.093 | 57 | 251 | 4.74 | 51 | 183.57 |
| 1100 | 0.0189 | 0.42 | 230.9 | 0.097 | 54 | 590 | 11.16 | 115 | 414.37 |

**Footnote.** Photon uptake was determined by quantifying light attenuation across the flat-panel photobioreactor cuvettes, comparing light transmitted through cuvettes filled with culture against medium-only controls, and converting the captured photon flux to a specific uptake rate using the measured cellular dry weight at each intensity. Values in the final column are rounded; the exact rates applied as model constraints.

**Table S7.** Enzyme lower-bound relaxations required to reach the measured growth rate under fixed measured enzyme abundances, for each light condition (27.5, 440, and 1100 µmol photons $m^{-2}$ $s^{-1}$). "Lower bound, original" is the enzyme usage bound set directly from the measured abundance; "Lower bound, relaxed" is the bound after relaxation; "Fold-change" is the ratio of relaxed to original bound; "µ after relaxation" is the growth rate attained in the model once that relaxation step was applied. Bounds are negative by the model's sign convention for enzyme usage reactions. Turnover numbers (Kcat) were not modified at this stage.

| Light condition (µmol photons $m^{-2}$ $s^{-1}$) | Enzyme (UniProt ID) | Lower bound, original (mmol $gDW^{-1}$ $h^{-1}$) | Lower bound, relaxed (mmol $gDW^{-1}$ $h^{-1}$) | Fold-change | µ after relaxation ($h^{-1}$) |
|---|---|---|---|---|---|
| 27.5 | P27178 | -0.09085 | -0.1363 | 1.5 | 0.0072 |
| 27.5 | P27178 | -0.09085 | -0.1817 | 2 | 0.0108 |
| 27.5 | P27178 | -0.09085 | -0.2726 | 3 | 0.01472 |
| 27.5 | P27178 | -0.09085 | -0.3634 | 4 | 0.0216 |
| 27.5 | P27181 | -0.336 | -0.504 | 1.5 | 0.02033 |
| 27.5 | P27182 | -0.01345 | -0.02017 | 1.5 | 0.00484 |
| 27.5 | P27182 | -0.01345 | -0.0269 | 2 | 0.00709 |
| 27.5 | P27182 | -0.01345 | -0.04035 | 3 | 0.0081 |
| 27.5 | P27182 | -0.01345 | -0.0538 | 4 | 0.01271 |
| 27.5 | P27182 | -0.01345 | -0.06725 | 5 | 0.0162 |
| 27.5 | P27182 | -0.01345 | -0.1345 | 10 | 0.02032 |
| 27.5 | P37294 | -0.01001 | -0.01502 | 1.5 | 0.00606 |

| Light condition (μmol photons $m^{-2}$ $s^{-1}$) | Enzyme (UniProt ID) | Lower bound, original (mmol $gDW^{-1}$ $h^{-1}$) | Lower bound, relaxed (mmol $gDW^{-1}$ $h^{-1}$) | Fold-change | μ after relaxation ($h^{-1}$) |
|---|---|---|---|---|---|
| 27.5 | P37294 | -0.01001 | -0.02002 | 2 | 0.00904 |
| 27.5 | P37294 | -0.01001 | -0.03003 | 3 | 0.01213 |
| 27.5 | P37294 | -0.01001 | -0.04004 | 4 | 0.01808 |
| 27.5 | P37294 | -0.01001 | -0.05005 | 5 | 0.02451 |
| 27.5 | P52965 | -0.004637 | -0.2318 | 50 | 0.00901 |
| 27.5 | P72652 | -0.00323 | -0.004846 | 1.5 | 0.01042 |
| 27.5 | P72652 | -0.00323 | -0.006461 | 2 | 0.01499 |
| 27.5 | P72652 | -0.00323 | -0.009691 | 3 | 0.02021 |
| 27.5 | P72663 | -0.008345 | -0.01252 | 1.5 | 0.02362 |
| 27.5 | P72842 | -0.01784 | -0.02677 | 1.5 | 0.00404 |
| 27.5 | P72842 | -0.01784 | -0.03569 | 2 | 0.00563 |
| 27.5 | P72842 | -0.01784 | -0.05353 | 3 | 0.00791 |
| 27.5 | P72842 | -0.01784 | -0.07138 | 4 | 0.01187 |
| 27.5 | P72842 | -0.01784 | -0.08922 | 5 | 0.01541 |
| 27.5 | P72842 | -0.01784 | -0.1784 | 10 | 0.01906 |
| 27.5 | P72934 | -0.01329 | -0.01994 | 1.5 | 0.01201 |
| 27.5 | P72934 | -0.01329 | -0.02659 | 2 | 0.01802 |
| 27.5 | P72934 | -0.01329 | -0.03988 | 3 | 0.02311 |
| 27.5 | P73426 | -0.01433 | -0.0215 | 1.5 | 0.01815 |
| 27.5 | P73448 | -0.01311 | -0.01967 | 1.5 | 0.00562 |
| 27.5 | P73448 | -0.01311 | -0.02622 | 2 | 0.00891 |
| 27.5 | P73448 | -0.01311 | -0.03933 | 3 | 0.01121 |
| 27.5 | P73448 | -0.01311 | -0.05244 | 4 | 0.01682 |
| 27.5 | P73448 | -0.01311 | -0.06556 | 5 | 0.02243 |
| 27.5 | P73449 | -0.05994 | -0.0899 | 1.5 | 0.00558 |
| 27.5 | P73449 | -0.05994 | -0.1199 | 2 | 0.00836 |
| 27.5 | P73449 | -0.05994 | -0.1798 | 3 | 0.01115 |
| 27.5 | P73449 | -0.05994 | -0.2397 | 4 | 0.01673 |
| 27.5 | P73449 | -0.05994 | -0.2997 | 5 | 0.0223 |

| Light condition (µmol photons $m^{-2}$ $s^{-1}$) | Enzyme (UniProt ID) | Lower bound, original (mmol $gDW^{-1}$ $h^{-1}$) | Lower bound, relaxed (mmol $gDW^{-1}$ $h^{-1}$) | Fold-change | µ after relaxation ($h^{-1}$) |
|---|---|---|---|---|---|
| 27.5 | P73450 | -0.1212 | -0.1818 | 1.5 | 0.00549 |
| 27.5 | P73450 | -0.1212 | -0.2424 | 2 | 0.00823 |
| 27.5 | P73450 | -0.1212 | -0.3636 | 3 | 0.01097 |
| 27.5 | P73450 | -0.1212 | -0.4848 | 4 | 0.01646 |
| 27.5 | P73450 | -0.1212 | -0.606 | 5 | 0.02195 |
| 27.5 | P73638 | -0.003773 | -0.00566 | 1.5 | 0.00911 |
| 27.5 | P73638 | -0.003773 | -0.007547 | 2 | 0.01418 |
| 27.5 | P73638 | -0.003773 | -0.01132 | 3 | 0.0181 |
| 27.5 | P73948 | -0.01056 | -0.01584 | 1.5 | 0.00897 |
| 27.5 | P73948 | -0.01056 | -0.02112 | 2 | 0.01356 |
| 27.5 | P73948 | -0.01056 | -0.03168 | 3 | 0.01799 |
| 27.5 | P74033 | -0.01626 | -0.02439 | 1.5 | 0.00981 |
| 27.5 | P74033 | -0.01626 | -0.06504 | 4 | 0.01281 |
| 27.5 | P74033 | -0.01626 | -0.0813 | 5 | 0.02127 |
| 27.5 | P74033 | -0.01626 | -0.1626 | 10 | 0.02478 |
| 27.5 | P74122 | -0.1208 | -0.1811 | 1.5 | 0.01574 |
| 27.5 | P74122 | -0.1208 | -0.2415 | 2 | 0.02328 |
| 27.5 | P74287 | -0.003144 | -0.004716 | 1.5 | 0.01583 |
| 27.5 | P74309 | -0.04761 | -0.09521 | 2 | 0.00375 |
| 27.5 | P74309 | -0.04761 | -0.1428 | 3 | 0.00601 |
| 27.5 | P74309 | -0.04761 | -0.1904 | 4 | 0.00809 |
| 27.5 | P74309 | -0.04761 | -0.238 | 5 | 0.01063 |
| 27.5 | P74309 | -0.04761 | -0.4761 | 10 | 0.01336 |
| 27.5 | P74309 | -0.04761 | -2.38 | 50 | 0.02524 |
| 27.5 | P74384 | -0.2089 | -0.3133 | 1.5 | 0.01962 |
| 27.5 | P74416 | -0.02431 | -0.03647 | 1.5 | 0.01617 |
| 27.5 | P74416 | -0.02431 | -0.04863 | 2 | 0.02374 |
| 27.5 | P74438 | -0.006251 | -0.009377 | 1.5 | 0.01782 |
| 27.5 | P74654 | -0.003811 | -0.005717 | 1.5 | 0.01874 |

| Light condition (µmol photons $m^{-2}$ $s^{-1}$) | Enzyme (UniProt ID) | Lower bound, original (mmol $gDW^{-1}$ $h^{-1}$) | Lower bound, relaxed (mmol $gDW^{-1}$ $h^{-1}$) | Fold-change | µ after relaxation ($h^{-1}$) |
|---|---|---|---|---|---|
| 27.5 | Q01903 | -0.002832 | -0.004249 | 1.5 | 0.01124 |
| 27.5 | Q01903 | -0.002832 | -0.005665 | 2 | 0.01692 |
| 27.5 | Q01903 | -0.002832 | -0.008497 | 3 | 0.02292 |
| 27.5 | Q55124 | -0.02041 | -0.03061 | 1.5 | 0.00749 |
| 27.5 | Q55124 | -0.02041 | -0.04082 | 2 | 0.0108 |
| 27.5 | Q55366 | -0.2909 | -0.4364 | 1.5 | 0.02114 |
| 27.5 | Q55664 | -5.533 | -276.6 | 50 | 0.0054 |
| 27.5 | Q55759 | -0.1645 | -0.2468 | 1.5 | 0.00802 |
| 27.5 | Q55759 | -0.1645 | -0.3291 | 2 | 0.01199 |
| 27.5 | Q55759 | -0.1645 | -0.4936 | 3 | 0.01604 |
| 27.5 | Q55759 | -0.1645 | -0.6582 | 4 | 0.02402 |
| 440 | P17253 | -1.029 | -1.543 | 1.5 | 0.07329 |
| 440 | P20388 | -0.01229 | -0.01844 | 1.5 | 0.08664 |
| 440 | P26527 | -5.317 | -7.975 | 1.5 | 0.08403 |
| 440 | P26533 | -0.1076 | -0.1614 | 1.5 | 0.018 |
| 440 | P26533 | -0.1076 | -0.2152 | 2 | 0.02748 |
| 440 | P26533 | -0.1076 | -0.3228 | 3 | 0.036 |
| 440 | P26533 | -0.1076 | -0.4304 | 4 | 0.05416 |
| 440 | P26533 | -0.1076 | -0.538 | 5 | 0.07221 |
| 440 | P26533 | -0.1076 | -1.076 | 10 | 0.0916 |
| 440 | P27178 | -0.07206 | -0.1081 | 1.5 | 0.00583 |
| 440 | P27178 | -0.07206 | -0.1441 | 2 | 0.00879 |
| 440 | P27178 | -0.07206 | -0.2162 | 3 | 0.01152 |
| 440 | P27178 | -0.07206 | -0.2882 | 4 | 0.01758 |
| 440 | P27178 | -0.07206 | -0.3603 | 5 | 0.02493 |
| 440 | P27178 | -0.07206 | -0.7206 | 10 | 0.02935 |
| 440 | P27178 | -0.07206 | -3.603 | 50 | 0.05703 |
| 440 | P27179 | -6.28 | -9.42 | 1.5 | 0.09597 |
| 440 | P27180 | -0.2106 | -0.3159 | 1.5 | 0.02554 |

| Light condition (μmol photons $m^{-2}$ $s^{-1}$) | Enzyme (UniProt ID) | Lower bound, original (mmol $gDW^{-1}$ $h^{-1}$) | Lower bound, relaxed (mmol $gDW^{-1}$ $h^{-1}$) | Fold-change | μ after relaxation ($h^{-1}$) |
|---|---|---|---|---|---|
| 440 | P27180 | -0.2106 | -0.4212 | 2 | 0.03831 |
| 440 | P27180 | -0.2106 | -0.6318 | 3 | 0.0516 |
| 440 | P27180 | -0.2106 | -0.8423 | 4 | 0.07806 |
| 440 | P27180 | -0.2106 | -1.053 | 5 | 0.10295 |
| 440 | P27181 | -0.2386 | -0.3579 | 1.5 | 0.01474 |
| 440 | P27181 | -0.2386 | -0.4772 | 2 | 0.0221 |
| 440 | P27181 | -0.2386 | -0.7159 | 3 | 0.02964 |
| 440 | P27181 | -0.2386 | -0.9545 | 4 | 0.04419 |
| 440 | P27181 | -0.2386 | -1.193 | 5 | 0.05932 |
| 440 | P27181 | -0.2386 | -2.386 | 10 | 0.07465 |
| 440 | P27182 | -0.01742 | -0.02613 | 1.5 | 0.0054 |
| 440 | P27182 | -0.01742 | -0.03484 | 2 | 0.00817 |
| 440 | P27182 | -0.01742 | -0.05226 | 3 | 0.01075 |
| 440 | P27182 | -0.01742 | -0.06967 | 4 | 0.01633 |
| 440 | P27182 | -0.01742 | -0.08709 | 5 | 0.02201 |
| 440 | P27182 | -0.01742 | -0.1742 | 10 | 0.02671 |
| 440 | P27182 | -0.01742 | -0.8709 | 50 | 0.05342 |
| 440 | P29107 | -1.813 | -2.719 | 1.5 | 0.08054 |
| 440 | P37294 | -0.009707 | -0.01456 | 1.5 | 0.00949 |
| 440 | P37294 | -0.009707 | -0.01941 | 2 | 0.01384 |
| 440 | P37294 | -0.009707 | -0.02912 | 3 | 0.01832 |
| 440 | P37294 | -0.009707 | -0.03883 | 4 | 0.02748 |
| 440 | P37294 | -0.009707 | -0.04854 | 5 | 0.03664 |
| 440 | P37294 | -0.009707 | -0.09707 | 10 | 0.04622 |
| 440 | P37294 | -0.009707 | -0.4854 | 50 | 0.09161 |
| 440 | P52965 | -0.003292 | -0.1646 | 50 | 0.0099 |
| 440 | P54261 | -0.02324 | -0.03486 | 1.5 | 0.04402 |
| 440 | P54261 | -0.02324 | -0.04648 | 2 | 0.06581 |
| 440 | P54261 | -0.02324 | -0.06973 | 3 | 0.08634 |

| Light condition (µmol photons m⁻² s⁻¹) | Enzyme (UniProt ID) | Lower bound, original (mmol gDW⁻¹ h⁻¹) | Lower bound, relaxed (mmol gDW⁻¹ h⁻¹) | Fold-change | µ after relaxation (h⁻¹) |
|---|---|---|---|---|---|
| 440 | P54386 | -0.02844 | -0.04266 | 1.5 | 0.08664 |
| 440 | P72652 | -0.003115 | -0.004673 | 1.5 | 0.01008 |
| 440 | P72652 | -0.003115 | -0.00623 | 2 | 0.01493 |
| 440 | P72652 | -0.003115 | -0.009345 | 3 | 0.02003 |
| 440 | P72652 | -0.003115 | -0.01246 | 4 | 0.02986 |
| 440 | P72652 | -0.003115 | -0.01558 | 5 | 0.03954 |
| 440 | P72652 | -0.003115 | -0.03115 | 10 | 0.05092 |
| 440 | P72652 | -0.003115 | -0.1558 | 50 | 0.09948 |
| 440 | P72663 | -0.00659 | -0.009885 | 1.5 | 0.03165 |
| 440 | P72663 | -0.00659 | -0.01318 | 2 | 0.04861 |
| 440 | P72663 | -0.00659 | -0.01977 | 3 | 0.06215 |
| 440 | P72663 | -0.00659 | -0.02636 | 4 | 0.09322 |
| 440 | P72794 | -0.006772 | -0.01016 | 1.5 | 0.03944 |
| 440 | P72794 | -0.006772 | -0.01354 | 2 | 0.0587 |
| 440 | P72794 | -0.006772 | -0.02032 | 3 | 0.08016 |
| 440 | P72797 | -0.3192 | -0.4788 | 1.5 | 0.04083 |
| 440 | P72797 | -0.3192 | -0.6384 | 2 | 0.06162 |
| 440 | P72797 | -0.3192 | -0.9577 | 3 | 0.08344 |
| 440 | P72842 | -0.01502 | -0.02253 | 1.5 | 0.00408 |
| 440 | P72842 | -0.01502 | -0.03004 | 2 | 0.00572 |
| 440 | P72842 | -0.01502 | -0.04506 | 3 | 0.00729 |
| 440 | P72842 | -0.01502 | -0.06007 | 4 | 0.01144 |
| 440 | P72842 | -0.01502 | -0.07509 | 5 | 0.01455 |
| 440 | P72842 | -0.01502 | -0.1502 | 10 | 0.01832 |
| 440 | P72842 | -0.01502 | -0.7509 | 50 | 0.03611 |
| 440 | P72934 | -0.009114 | -0.01367 | 1.5 | 0.00858 |
| 440 | P72934 | -0.009114 | -0.01823 | 2 | 0.01225 |
| 440 | P72934 | -0.009114 | -0.02734 | 3 | 0.01575 |
| 440 | P72934 | -0.009114 | -0.03646 | 4 | 0.02222 |

| Light condition (μmol photons $m^{-2}$ $s^{-1}$) | Enzyme (UniProt ID) | Lower bound, original (mmol $gDW^{-1}$ $h^{-1}$) | Lower bound, relaxed (mmol $gDW^{-1}$ $h^{-1}$) | Fold-change | μ after relaxation ($h^{-1}$) |
|---|---|---|---|---|---|
| 440 | P72934 | -0.009114 | -0.04557 | 5 | 0.02859 |
| 440 | P72934 | -0.009114 | -0.09114 | 10 | 0.0354 |
| 440 | P72934 | -0.009114 | -0.4557 | 50 | 0.06855 |
| 440 | P73016 | -0.06766 | -0.1015 | 1.5 | 0.0615 |
| 440 | P73016 | -0.06766 | -0.1353 | 2 | 0.09243 |
| 440 | P73053 | -0.02723 | -0.04084 | 1.5 | 0.03443 |
| 440 | P73053 | -0.02723 | -0.05446 | 2 | 0.05105 |
| 440 | P73053 | -0.02723 | -0.08169 | 3 | 0.0681 |
| 440 | P73053 | -0.02723 | -0.1089 | 4 | 0.10212 |
| 440 | P73071 | -0.005283 | -0.007924 | 1.5 | 0.08838 |
| 440 | P73133 | -0.1539 | -0.2308 | 1.5 | 0.05972 |
| 440 | P73133 | -0.1539 | -0.3078 | 2 | 0.08345 |
| 440 | P73426 | -0.0105 | -0.01575 | 1.5 | 0.0224 |
| 440 | P73426 | -0.0105 | -0.021 | 2 | 0.0336 |
| 440 | P73426 | -0.0105 | -0.0315 | 3 | 0.04462 |
| 440 | P73426 | -0.0105 | -0.042 | 4 | 0.06677 |
| 440 | P73426 | -0.0105 | -0.05249 | 5 | 0.08966 |
| 440 | P73448 | -0.01028 | -0.01543 | 1.5 | 0.00533 |
| 440 | P73448 | -0.01028 | -0.02057 | 2 | 0.00799 |
| 440 | P73448 | -0.01028 | -0.03085 | 3 | 0.01065 |
| 440 | P73448 | -0.01028 | -0.04113 | 4 | 0.01598 |
| 440 | P73448 | -0.01028 | -0.05142 | 5 | 0.0213 |
| 440 | P73448 | -0.01028 | -0.1028 | 10 | 0.02639 |
| 440 | P73448 | -0.01028 | -0.5142 | 50 | 0.05325 |
| 440 | P73449 | -0.06003 | -0.09004 | 1.5 | 0.00672 |
| 440 | P73449 | -0.06003 | -0.1201 | 2 | 0.00992 |
| 440 | P73449 | -0.06003 | -0.1801 | 3 | 0.01335 |
| 440 | P73449 | -0.06003 | -0.2401 | 4 | 0.01981 |
| 440 | P73449 | -0.06003 | -0.3001 | 5 | 0.02663 |

| Light condition (μmol photons $m^{-2}$ $s^{-1}$) | Enzyme (UniProt ID) | Lower bound, original (mmol $gDW^{-1}$ $h^{-1}$) | Lower bound, relaxed (mmol $gDW^{-1}$ $h^{-1}$) | Fold-change | μ after relaxation ($h^{-1}$) |
|---|---|---|---|---|---|
| 440 | P73449 | -0.06003 | -0.6003 | 10 | 0.03314 |
| 440 | P73449 | -0.06003 | -3.001 | 50 | 0.06629 |
| 440 | P73450 | -0.1256 | -0.1884 | 1.5 | 0.00687 |
| 440 | P73450 | -0.1256 | -0.2512 | 2 | 0.01038 |
| 440 | P73450 | -0.1256 | -0.3768 | 3 | 0.01374 |
| 440 | P73450 | -0.1256 | -0.5023 | 4 | 0.02042 |
| 440 | P73450 | -0.1256 | -0.6279 | 5 | 0.02697 |
| 440 | P73450 | -0.1256 | -1.256 | 10 | 0.03395 |
| 440 | P73450 | -0.1256 | -6.279 | 50 | 0.06789 |
| 440 | P73452 | -0.6903 | -1.035 | 1.5 | 0.05673 |
| 440 | P73452 | -0.6903 | -1.381 | 2 | 0.08555 |
| 440 | P73617 | -0.0157 | -0.02355 | 1.5 | 0.1044 |
| 440 | P73638 | -0.002756 | -0.004134 | 1.5 | 0.00716 |
| 440 | P73638 | -0.002756 | -0.005512 | 2 | 0.0105 |
| 440 | P73638 | -0.002756 | -0.008268 | 3 | 0.01433 |
| 440 | P73638 | -0.002756 | -0.01102 | 4 | 0.021 |
| 440 | P73638 | -0.002756 | -0.01378 | 5 | 0.02852 |
| 440 | P73638 | -0.002756 | -0.02756 | 10 | 0.03498 |
| 440 | P73638 | -0.002756 | -0.1378 | 50 | 0.0708 |
| 440 | P73761 | -0.006304 | -0.009456 | 1.5 | 0.05356 |
| 440 | P73761 | -0.006304 | -0.01261 | 2 | 0.08034 |
| 440 | P73948 | -0.005086 | -0.007629 | 1.5 | 0.00438 |
| 440 | P73948 | -0.005086 | -0.01017 | 2 | 0.00619 |
| 440 | P73948 | -0.005086 | -0.01526 | 3 | 0.00831 |
| 440 | P73948 | -0.005086 | -0.02035 | 4 | 0.01237 |
| 440 | P73948 | -0.005086 | -0.02543 | 5 | 0.01715 |
| 440 | P73948 | -0.005086 | -0.05086 | 10 | 0.02073 |
| 440 | P73948 | -0.005086 | -0.2543 | 50 | 0.04277 |
| 440 | P74033 | -0.008261 | -0.01239 | 1.5 | 0.00525 |

| Light condition (μmol photons $m^{-2}$ $s^{-1}$) | Enzyme (UniProt ID) | Lower bound, original (mmol $gDW^{-1}$ $h^{-1}$) | Lower bound, relaxed (mmol $gDW^{-1}$ $h^{-1}$) | Fold-change | μ after relaxation ($h^{-1}$) |
|---|---|---|---|---|---|
| 440 | P74033 | -0.008261 | -0.01652 | 2 | 0.00633 |
| 440 | P74033 | -0.008261 | -0.02478 | 3 | 0.00916 |
| 440 | P74033 | -0.008261 | -0.03304 | 4 | 0.01266 |
| 440 | P74033 | -0.008261 | -0.0413 | 5 | 0.01798 |
| 440 | P74033 | -0.008261 | -0.08261 | 10 | 0.02287 |
| 440 | P74033 | -0.008261 | -0.413 | 50 | 0.0525 |
| 440 | P74122 | -0.1017 | -0.1526 | 1.5 | 0.03257 |
| 440 | P74122 | -0.1017 | -0.2035 | 2 | 0.04926 |
| 440 | P74122 | -0.1017 | -0.3052 | 3 | 0.0517 |
| 440 | P74122 | -0.1017 | -0.407 | 4 | 0.08065 |
| 440 | P74122 | -0.1017 | -0.5087 | 5 | 0.08497 |
| 440 | P74122 | -0.1017 | -1.017 | 10 | 0.09697 |
| 440 | P74208 | -0.0681 | -0.1022 | 1.5 | 0.08164 |
| 440 | P74240 | -0.02675 | -0.04012 | 1.5 | 0.06329 |
| 440 | P74240 | -0.02675 | -0.0535 | 2 | 0.0945 |
| 440 | P74287 | -0.002126 | -0.004253 | 2 | 0.02532 |
| 440 | P74287 | -0.002126 | -0.006379 | 3 | 0.0258 |
| 440 | P74287 | -0.002126 | -0.01063 | 5 | 0.03903 |
| 440 | P74309 | -0.0285 | -0.0855 | 3 | 0.0036 |
| 440 | P74309 | -0.0285 | -0.114 | 4 | 0.00521 |
| 440 | P74309 | -0.0285 | -0.1425 | 5 | 0.00612 |
| 440 | P74309 | -0.0285 | -0.285 | 10 | 0.00737 |
| 440 | P74309 | -0.0285 | -1.425 | 50 | 0.01468 |
| 440 | P74309 | -0.0285 | -28.5 | 1000 | 0.0615 |
| 440 | P74384 | -0.1449 | -0.2173 | 1.5 | 0.01344 |
| 440 | P74384 | -0.1449 | -0.2898 | 2 | 0.02016 |
| 440 | P74384 | -0.1449 | -0.4347 | 3 | 0.02688 |
| 440 | P74384 | -0.1449 | -0.5795 | 4 | 0.04033 |
| 440 | P74384 | -0.1449 | -0.7244 | 5 | 0.05344 |

| Light condition (μmol photons $m^{-2}$ $s^{-1}$) | Enzyme (UniProt ID) | Lower bound, original (mmol $gDW^{-1}$ $h^{-1}$) | Lower bound, relaxed (mmol $gDW^{-1}$ $h^{-1}$) | Fold-change | μ after relaxation ($h^{-1}$) |
|---|---|---|---|---|---|
| 440 | P74384 | -0.1449 | -1.449 | 10 | 0.06716 |
| 440 | P74416 | -0.06182 | -0.09273 | 1.5 | 0.04479 |
| 440 | P74416 | -0.06182 | -0.1236 | 2 | 0.06597 |
| 440 | P74416 | -0.06182 | -0.1855 | 3 | 0.08752 |
| 440 | P74438 | -0.002881 | -0.005763 | 2 | 0.00788 |
| 440 | P74438 | -0.002881 | -0.008644 | 3 | 0.01487 |
| 440 | P74438 | -0.002881 | -0.01153 | 4 | 0.0672 |
| 440 | P74625 | -0.07782 | -0.1167 | 1.5 | 0.05854 |
| 440 | P74625 | -0.07782 | -0.1556 | 2 | 0.087 |
| 440 | P74638 | -0.02949 | -0.04423 | 1.5 | 0.03664 |
| 440 | P74638 | -0.02949 | -0.05898 | 2 | 0.05443 |
| 440 | P74638 | -0.02949 | -0.08846 | 3 | 0.07243 |
| 440 | P74654 | -0.003911 | -0.005866 | 1.5 | 0.01954 |
| 440 | P74654 | -0.003911 | -0.007821 | 2 | 0.02616 |
| 440 | P74654 | -0.003911 | -0.01173 | 3 | 0.03582 |
| 440 | P74654 | -0.003911 | -0.01564 | 4 | 0.05768 |
| 440 | P74654 | -0.003911 | -0.01955 | 5 | 0.07191 |
| 440 | P74724 | -0.02017 | -0.03025 | 1.5 | 0.06049 |
| 440 | P74724 | -0.02017 | -0.04034 | 2 | 0.08989 |
| 440 | P74755 | -0.0036 | -0.0054 | 1.5 | 0.06665 |
| 440 | P74755 | -0.0036 | -0.007199 | 2 | 0.10034 |
| 440 | Q01903 | -0.003139 | -0.004709 | 1.5 | 0.01568 |
| 440 | Q01903 | -0.003139 | -0.006278 | 2 | 0.02279 |
| 440 | Q01903 | -0.003139 | -0.009418 | 3 | 0.03081 |
| 440 | Q01903 | -0.003139 | -0.01256 | 4 | 0.0458 |
| 440 | Q01903 | -0.003139 | -0.0157 | 5 | 0.05977 |
| 440 | Q01903 | -0.003139 | -0.03139 | 10 | 0.07602 |
| 440 | Q55120 | -0.01792 | -0.02688 | 1.5 | 0.09807 |
| 440 | Q55124 | -0.0152 | -0.02279 | 1.5 | 0.0072 |

| Light condition (μmol photons $m^{-2}$ $s^{-1}$) | Enzyme (UniProt ID) | Lower bound, original (mmol $gDW^{-1}$ $h^{-1}$) | Lower bound, relaxed (mmol $gDW^{-1}$ $h^{-1}$) | Fold-change | μ after relaxation ($h^{-1}$) |
|---|---|---|---|---|---|
| 440 | Q55124 | -0.0152 | -0.03039 | 2 | 0.0108 |
| 440 | Q55124 | -0.0152 | -0.04559 | 3 | 0.0144 |
| 440 | Q55124 | -0.0152 | -0.06079 | 4 | 0.01979 |
| 440 | Q55124 | -0.0152 | -0.07598 | 5 | 0.03298 |
| 440 | Q55124 | -0.0152 | -0.152 | 10 | 0.03596 |
| 440 | Q55160 | -0.04826 | -0.07239 | 1.5 | 0.08687 |
| 440 | Q55366 | -0.2183 | -0.3275 | 1.5 | 0.01899 |
| 440 | Q55366 | -0.2183 | -0.4366 | 2 | 0.02759 |
| 440 | Q55366 | -0.2183 | -0.655 | 3 | 0.03829 |
| 440 | Q55366 | -0.2183 | -0.8733 | 4 | 0.05665 |
| 440 | Q55366 | -0.2183 | -1.092 | 5 | 0.07338 |
| 440 | Q55366 | -0.2183 | -2.183 | 10 | 0.09224 |
| 440 | Q55406 | -0.01409 | -0.02114 | 1.5 | 0.09973 |
| 440 | Q55422 | -0.03579 | -0.05368 | 1.5 | 0.04076 |
| 440 | Q55422 | -0.03579 | -0.07157 | 2 | 0.06111 |
| 440 | Q55422 | -0.03579 | -0.1074 | 3 | 0.08136 |
| 440 | Q55515 | -0.001916 | -0.002874 | 1.5 | 0.02211 |
| 440 | Q55515 | -0.001916 | -0.003832 | 2 | 0.03332 |
| 440 | Q55515 | -0.001916 | -0.005748 | 3 | 0.04443 |
| 440 | Q55515 | -0.001916 | -0.007664 | 4 | 0.06649 |
| 440 | Q55515 | -0.001916 | -0.00958 | 5 | 0.08886 |
| 440 | Q55574 | -0.0382 | -0.05729 | 1.5 | 0.02855 |
| 440 | Q55574 | -0.0382 | -0.07639 | 2 | 0.04391 |
| 440 | Q55574 | -0.0382 | -0.1146 | 3 | 0.05758 |
| 440 | Q55574 | -0.0382 | -0.1528 | 4 | 0.08558 |
| 440 | Q55626 | -0.002291 | -0.003436 | 1.5 | 0.02625 |
| 440 | Q55626 | -0.002291 | -0.004582 | 2 | 0.04006 |
| 440 | Q55626 | -0.002291 | -0.006872 | 3 | 0.05209 |
| 440 | Q55626 | -0.002291 | -0.009163 | 4 | 0.07659 |

| Light condition (μmol photons $m^{-2}$ $s^{-1}$) | Enzyme (UniProt ID) | Lower bound, original (mmol $gDW^{-1}$ $h^{-1}$) | Lower bound, relaxed (mmol $gDW^{-1}$ $h^{-1}$) | Fold-change | μ after relaxation ($h^{-1}$) |
|---|---|---|---|---|---|
| 440 | Q55626 | -0.002291 | -0.01145 | 5 | 0.10184 |
| 440 | Q55643 | -0.01524 | -0.02286 | 1.5 | 0.05497 |
| 440 | Q55643 | -0.01524 | -0.03048 | 2 | 0.08177 |
| 440 | Q55759 | -0.1383 | -0.2074 | 1.5 | 0.0066 |
| 440 | Q55759 | -0.1383 | -0.2765 | 2 | 0.00984 |
| 440 | Q55759 | -0.1383 | -0.4148 | 3 | 0.01319 |
| 440 | Q55759 | -0.1383 | -0.5531 | 4 | 0.01919 |
| 440 | Q55759 | -0.1383 | -0.6913 | 5 | 0.02553 |
| 440 | Q55759 | -0.1383 | -1.383 | 10 | 0.03241 |
| 440 | Q55759 | -0.1383 | -6.913 | 50 | 0.06474 |
| 440 | Q57417 | -0.04417 | -0.06625 | 1.5 | 0.05393 |
| 440 | Q57417 | -0.04417 | -0.08834 | 2 | 0.08051 |
| 1100 | P17253 | -1.045 | -1.568 | 1.5 | 0.07427 |
| 1100 | P20388 | -0.008393 | -0.01259 | 1.5 | 0.06253 |
| 1100 | P20388 | -0.008393 | -0.01679 | 2 | 0.09264 |
| 1100 | P26527 | -5.382 | -8.073 | 1.5 | 0.08503 |
| 1100 | P26533 | -0.1309 | -0.1963 | 1.5 | 0.02213 |
| 1100 | P26533 | -0.1309 | -0.2618 | 2 | 0.03318 |
| 1100 | P26533 | -0.1309 | -0.3926 | 3 | 0.04326 |
| 1100 | P26533 | -0.1309 | -0.5235 | 4 | 0.06613 |
| 1100 | P26533 | -0.1309 | -0.6544 | 5 | 0.0882 |
| 1100 | P27178 | -0.05282 | -0.07924 | 1.5 | 0.00393 |
| 1100 | P27178 | -0.05282 | -0.1056 | 2 | 0.0063 |
| 1100 | P27178 | -0.05282 | -0.1585 | 3 | 0.0087 |
| 1100 | P27178 | -0.05282 | -0.2113 | 4 | 0.01279 |
| 1100 | P27178 | -0.05282 | -0.2641 | 5 | 0.01688 |
| 1100 | P27178 | -0.05282 | -0.5282 | 10 | 0.02139 |
| 1100 | P27178 | -0.05282 | -2.641 | 50 | 0.04104 |
| 1100 | P27180 | -0.2159 | -0.3239 | 1.5 | 0.02799 |

| Light condition (μmol photons $m^{-2}$ $s^{-1}$) | Enzyme (UniProt ID) | Lower bound, original (mmol $gDW^{-1}$ $h^{-1}$) | Lower bound, relaxed (mmol $gDW^{-1}$ $h^{-1}$) | Fold-change | μ after relaxation ($h^{-1}$) |
|---|---|---|---|---|---|
| 1100 | P27180 | -0.2159 | -0.4318 | 2 | 0.03912 |
| 1100 | P27180 | -0.2159 | -0.6478 | 3 | 0.05035 |
| 1100 | P27180 | -0.2159 | -0.8637 | 4 | 0.08173 |
| 1100 | P27181 | -0.2388 | -0.3582 | 1.5 | 0.01538 |
| 1100 | P27181 | -0.2388 | -0.4777 | 2 | 0.02251 |
| 1100 | P27181 | -0.2388 | -0.7165 | 3 | 0.02974 |
| 1100 | P27181 | -0.2388 | -0.9553 | 4 | 0.04497 |
| 1100 | P27181 | -0.2388 | -1.194 | 5 | 0.05948 |
| 1100 | P27181 | -0.2388 | -2.388 | 10 | 0.07392 |
| 1100 | P27182 | -0.02138 | -0.03207 | 1.5 | 0.00671 |
| 1100 | P27182 | -0.02138 | -0.04276 | 2 | 0.01001 |
| 1100 | P27182 | -0.02138 | -0.06414 | 3 | 0.01371 |
| 1100 | P27182 | -0.02138 | -0.08552 | 4 | 0.02052 |
| 1100 | P27182 | -0.02138 | -0.1069 | 5 | 0.0263 |
| 1100 | P27182 | -0.02138 | -0.2138 | 10 | 0.03294 |
| 1100 | P27182 | -0.02138 | -1.069 | 50 | 0.06589 |
| 1100 | P29107 | -1.675 | -2.512 | 1.5 | 0.07889 |
| 1100 | P37294 | -0.01124 | -0.01686 | 1.5 | 0.01047 |
| 1100 | P37294 | -0.01124 | -0.02248 | 2 | 0.01552 |
| 1100 | P37294 | -0.01124 | -0.03372 | 3 | 0.02105 |
| 1100 | P37294 | -0.01124 | -0.04497 | 4 | 0.03148 |
| 1100 | P37294 | -0.01124 | -0.05621 | 5 | 0.04113 |
| 1100 | P37294 | -0.01124 | -0.1124 | 10 | 0.05474 |
| 1100 | P52965 | -0.004318 | -0.4318 | 100 | 0.0415 |
| 1100 | P52965 | -0.004318 | -2.159 | 500 | 0.08337 |
| 1100 | P54261 | -0.01763 | -0.02645 | 1.5 | 0.03334 |
| 1100 | P54261 | -0.01763 | -0.03526 | 2 | 0.04982 |
| 1100 | P54261 | -0.01763 | -0.0529 | 3 | 0.06667 |
| 1100 | P72652 | -0.003526 | -0.005289 | 1.5 | 0.01153 |

| Light condition (μmol photons $m^{-2}$ $s^{-1}$) | Enzyme (UniProt ID) | Lower bound, original (mmol $gDW^{-1}$ $h^{-1}$) | Lower bound, relaxed (mmol $gDW^{-1}$ $h^{-1}$) | Fold-change | μ after relaxation ($h^{-1}$) |
|---|---|---|---|---|---|
| 1100 | P72652 | -0.003526 | -0.007052 | 2 | 0.01708 |
| 1100 | P72652 | -0.003526 | -0.01058 | 3 | 0.02307 |
| 1100 | P72652 | -0.003526 | -0.0141 | 4 | 0.03417 |
| 1100 | P72652 | -0.003526 | -0.01763 | 5 | 0.04555 |
| 1100 | P72652 | -0.003526 | -0.03526 | 10 | 0.05668 |
| 1100 | P72663 | -0.006251 | -0.009377 | 1.5 | 0.03269 |
| 1100 | P72663 | -0.006251 | -0.0125 | 2 | 0.04904 |
| 1100 | P72663 | -0.006251 | -0.01875 | 3 | 0.0643 |
| 1100 | P72794 | -0.009427 | -0.01414 | 1.5 | 0.05729 |
| 1100 | P72794 | -0.009427 | -0.01885 | 2 | 0.08651 |
| 1100 | P72797 | -0.3921 | -0.5882 | 1.5 | 0.05125 |
| 1100 | P72797 | -0.3921 | -0.7842 | 2 | 0.07639 |
| 1100 | P72842 | -0.01406 | -0.02109 | 1.5 | 0.00385 |
| 1100 | P72842 | -0.01406 | -0.02812 | 2 | 0.00537 |
| 1100 | P72842 | -0.01406 | -0.04218 | 3 | 0.00712 |
| 1100 | P72842 | -0.01406 | -0.05625 | 4 | 0.01069 |
| 1100 | P72842 | -0.01406 | -0.07031 | 5 | 0.01426 |
| 1100 | P72842 | -0.01406 | -0.1406 | 10 | 0.01784 |
| 1100 | P72842 | -0.01406 | -0.7031 | 50 | 0.03565 |
| 1100 | P72934 | -0.01196 | -0.01795 | 1.5 | 0.01006 |
| 1100 | P72934 | -0.01196 | -0.02393 | 2 | 0.01475 |
| 1100 | P72934 | -0.01196 | -0.03589 | 3 | 0.0197 |
| 1100 | P72934 | -0.01196 | -0.04785 | 4 | 0.02834 |
| 1100 | P72934 | -0.01196 | -0.05982 | 5 | 0.03693 |
| 1100 | P72934 | -0.01196 | -0.1196 | 10 | 0.0482 |
| 1100 | P72934 | -0.01196 | -0.5982 | 50 | 0.0917 |
| 1100 | P73016 | -0.05915 | -0.08872 | 1.5 | 0.05605 |
| 1100 | P73016 | -0.05915 | -0.1183 | 2 | 0.08392 |
| 1100 | P73053 | -0.03019 | -0.04529 | 1.5 | 0.04052 |

| Light condition (μmol photons $m^{-2}$ $s^{-1}$) | Enzyme (UniProt ID) | Lower bound, original (mmol $gDW^{-1}$ $h^{-1}$) | Lower bound, relaxed (mmol $gDW^{-1}$ $h^{-1}$) | Fold-change | μ after relaxation ($h^{-1}$) |
|---|---|---|---|---|---|
| 1100 | P73053 | -0.03019 | -0.06039 | 2 | 0.05877 |
| 1100 | P73053 | -0.03019 | -0.09058 | 3 | 0.07859 |
| 1100 | P73071 | -0.004069 | -0.006104 | 1.5 | 0.06488 |
| 1100 | P73133 | -0.2071 | -0.3107 | 1.5 | 0.08429 |
| 1100 | P73426 | -0.01218 | -0.01827 | 1.5 | 0.02852 |
| 1100 | P73426 | -0.01218 | -0.02436 | 2 | 0.04321 |
| 1100 | P73426 | -0.01218 | -0.03655 | 3 | 0.05696 |
| 1100 | P73426 | -0.01218 | -0.04873 | 4 | 0.08542 |
| 1100 | P73448 | -0.01181 | -0.01772 | 1.5 | 0.0065 |
| 1100 | P73448 | -0.01181 | -0.02362 | 2 | 0.0096 |
| 1100 | P73448 | -0.01181 | -0.03544 | 3 | 0.01263 |
| 1100 | P73448 | -0.01181 | -0.04725 | 4 | 0.0192 |
| 1100 | P73448 | -0.01181 | -0.05906 | 5 | 0.02563 |
| 1100 | P73448 | -0.01181 | -0.1181 | 10 | 0.03189 |
| 1100 | P73448 | -0.01181 | -0.5906 | 50 | 0.06427 |
| 1100 | P73449 | -0.05183 | -0.07775 | 1.5 | 0.00629 |
| 1100 | P73449 | -0.05183 | -0.1037 | 2 | 0.00945 |
| 1100 | P73449 | -0.05183 | -0.1555 | 3 | 0.01222 |
| 1100 | P73449 | -0.05183 | -0.2073 | 4 | 0.01825 |
| 1100 | P73449 | -0.05183 | -0.2592 | 5 | 0.02452 |
| 1100 | P73449 | -0.05183 | -0.5183 | 10 | 0.02998 |
| 1100 | P73449 | -0.05183 | -2.592 | 50 | 0.05996 |
| 1100 | P73450 | -0.1276 | -0.1914 | 1.5 | 0.00733 |
| 1100 | P73450 | -0.1276 | -0.2552 | 2 | 0.01121 |
| 1100 | P73450 | -0.1276 | -0.3828 | 3 | 0.0144 |
| 1100 | P73450 | -0.1276 | -0.5104 | 4 | 0.02196 |
| 1100 | P73450 | -0.1276 | -0.638 | 5 | 0.02951 |
| 1100 | P73450 | -0.1276 | -1.276 | 10 | 0.03677 |
| 1100 | P73450 | -0.1276 | -6.38 | 50 | 0.07244 |

| Light condition (μmol photons $m^{-2}$ $s^{-1}$) | Enzyme (UniProt ID) | Lower bound, original (mmol $gDW^{-1}$ $h^{-1}$) | Lower bound, relaxed (mmol $gDW^{-1}$ $h^{-1}$) | Fold-change | μ after relaxation ($h^{-1}$) |
|---|---|---|---|---|---|
| 1100 | P73452 | -0.6558 | -0.9836 | 1.5 | 0.05662 |
| 1100 | P73452 | -0.6558 | -1.312 | 2 | 0.08486 |
| 1100 | P73638 | -0.003295 | -0.004942 | 1.5 | 0.00842 |
| 1100 | P73638 | -0.003295 | -0.00659 | 2 | 0.01259 |
| 1100 | P73638 | -0.003295 | -0.009884 | 3 | 0.01674 |
| 1100 | P73638 | -0.003295 | -0.01318 | 4 | 0.02519 |
| 1100 | P73638 | -0.003295 | -0.01647 | 5 | 0.03283 |
| 1100 | P73638 | -0.003295 | -0.03295 | 10 | 0.04251 |
| 1100 | P73638 | -0.003295 | -0.1647 | 50 | 0.08381 |
| 1100 | P73761 | -0.007298 | -0.01095 | 1.5 | 0.06456 |
| 1100 | P73906 | -0.003117 | -0.004675 | 1.5 | 0.07579 |
| 1100 | P73948 | -0.003137 | -0.004706 | 1.5 | 0.0027 |
| 1100 | P73948 | -0.003137 | -0.006275 | 2 | 0.00403 |
| 1100 | P73948 | -0.003137 | -0.009412 | 3 | 0.00524 |
| 1100 | P73948 | -0.003137 | -0.01255 | 4 | 0.00805 |
| 1100 | P73948 | -0.003137 | -0.01569 | 5 | 0.01048 |
| 1100 | P73948 | -0.003137 | -0.03137 | 10 | 0.01313 |
| 1100 | P73948 | -0.003137 | -0.1569 | 50 | 0.02626 |
| 1100 | P73953 | -0.05345 | -0.1604 | 3 | 0.04169 |
| 1100 | P73953 | -0.05345 | -0.5345 | 10 | 0.05571 |
| 1100 | P73953 | -0.05345 | -2.673 | 50 | 0.05619 |
| 1100 | P74033 | -0.01112 | -0.01668 | 1.5 | 0.00769 |
| 1100 | P74033 | -0.01112 | -0.02224 | 2 | 0.00892 |
| 1100 | P74033 | -0.01112 | -0.03336 | 3 | 0.01026 |
| 1100 | P74033 | -0.01112 | -0.04448 | 4 | 0.01309 |
| 1100 | P74033 | -0.01112 | -0.0556 | 5 | 0.01635 |
| 1100 | P74033 | -0.01112 | -0.1112 | 10 | 0.01879 |
| 1100 | P74033 | -0.01112 | -0.556 | 50 | 0.03375 |
| 1100 | P74122 | -0.1162 | -0.1744 | 1.5 | 0.03342 |

| Light condition (μmol photons $m^{-2}$ $s^{-1}$) | Enzyme (UniProt ID) | Lower bound, original (mmol $gDW^{-1}$ $h^{-1}$) | Lower bound, relaxed (mmol $gDW^{-1}$ $h^{-1}$) | Fold-change | μ after relaxation ($h^{-1}$) |
|---|---|---|---|---|---|
| 1100 | P74122 | -0.1162 | -0.2325 | 2 | 0.03714 |
| 1100 | P74208 | -0.07103 | -0.1066 | 1.5 | 0.08881 |
| 1100 | P74240 | -0.03678 | -0.05517 | 1.5 | 0.08994 |
| 1100 | P74287 | -0.002146 | -0.003219 | 1.5 | 0.05869 |
| 1100 | P74309 | -0.02619 | -0.05237 | 2 | 0.00262 |
| 1100 | P74309 | -0.02619 | -0.07856 | 3 | 0.00349 |
| 1100 | P74309 | -0.02619 | -0.1047 | 4 | 0.00524 |
| 1100 | P74309 | -0.02619 | -0.1309 | 5 | 0.00562 |
| 1100 | P74309 | -0.02619 | -0.2619 | 10 | 0.00713 |
| 1100 | P74309 | -0.02619 | -1.309 | 50 | 0.01398 |
| 1100 | P74309 | -0.02619 | -2.619 | 100 | 0.06834 |
| 1100 | P74384 | -0.1737 | -0.2605 | 1.5 | 0.01653 |
| 1100 | P74384 | -0.1737 | -0.3473 | 2 | 0.02488 |
| 1100 | P74384 | -0.1737 | -0.521 | 3 | 0.03279 |
| 1100 | P74384 | -0.1737 | -0.6947 | 4 | 0.04954 |
| 1100 | P74384 | -0.1737 | -0.8684 | 5 | 0.06565 |
| 1100 | P74384 | -0.1737 | -1.737 | 10 | 0.08324 |
| 1100 | P74416 | -0.09581 | -0.1437 | 1.5 | 0.06878 |
| 1100 | P74438 | -0.002439 | -0.003659 | 1.5 | 0.00985 |
| 1100 | P74438 | -0.002439 | -0.004878 | 2 | 0.01889 |
| 1100 | P74438 | -0.002439 | -0.007317 | 3 | 0.04585 |
| 1100 | P74625 | -0.05462 | -0.08193 | 1.5 | 0.04169 |
| 1100 | P74625 | -0.05462 | -0.1092 | 2 | 0.06297 |
| 1100 | P74625 | -0.05462 | -0.1639 | 3 | 0.0835 |
| 1100 | P74638 | -0.03001 | -0.04502 | 1.5 | 0.03845 |
| 1100 | P74638 | -0.03001 | -0.06002 | 2 | 0.05845 |
| 1100 | P74638 | -0.03001 | -0.09004 | 3 | 0.07689 |
| 1100 | P74654 | -0.007363 | -0.01473 | 2 | 0.04062 |
| 1100 | P74654 | -0.007363 | -0.02209 | 3 | 0.06696 |

| Light condition (μmol photons $m^{-2}$ $s^{-1}$) | Enzyme (UniProt ID) | Lower bound, original (mmol $gDW^{-1}$ $h^{-1}$) | Lower bound, relaxed (mmol $gDW^{-1}$ $h^{-1}$) | Fold-change | μ after relaxation ($h^{-1}$) |
|---|---|---|---|---|---|
| 1100 | P74724 | -0.01642 | -0.02463 | 1.5 | 0.04983 |
| 1100 | P74724 | -0.01642 | -0.03283 | 2 | 0.07553 |
| 1100 | P74755 | -0.003712 | -0.005568 | 1.5 | 0.07377 |
| 1100 | Q01903 | -0.0092 | -0.0138 | 1.5 | 0.04637 |
| 1100 | Q01903 | -0.0092 | -0.0184 | 2 | 0.06969 |
| 1100 | Q01903 | -0.0092 | -0.0276 | 3 | 0.0919 |
| 1100 | Q55124 | -0.01383 | -0.02074 | 1.5 | 0.00595 |
| 1100 | Q55124 | -0.01383 | -0.02765 | 2 | 0.03213 |
| 1100 | Q55366 | -0.1949 | -0.2924 | 1.5 | 0.01747 |
| 1100 | Q55366 | -0.1949 | -0.3899 | 2 | 0.02618 |
| 1100 | Q55366 | -0.1949 | -0.5848 | 3 | 0.03495 |
| 1100 | Q55366 | -0.1949 | -0.7798 | 4 | 0.0513 |
| 1100 | Q55366 | -0.1949 | -0.9747 | 5 | 0.06867 |
| 1100 | Q55366 | -0.1949 | -1.949 | 10 | 0.08543 |
| 1100 | Q55406 | -0.009499 | -0.01425 | 1.5 | 0.07129 |
| 1100 | Q55415 | -0.01141 | -5.707 | 500 | 0.0058 |
| 1100 | Q55415 | -0.01141 | -11.41 | 1000 | 0.00827 |
| 1100 | Q55415 | -0.01141 | -57.07 | 5000 | 0.0124 |
| 1100 | Q55422 | -0.03749 | -0.05624 | 1.5 | 0.04346 |
| 1100 | Q55422 | -0.03749 | -0.07499 | 2 | 0.06538 |
| 1100 | Q55422 | -0.03749 | -0.1125 | 3 | 0.08768 |
| 1100 | Q55574 | -0.03864 | -0.05796 | 1.5 | 0.03076 |
| 1100 | Q55574 | -0.03864 | -0.07727 | 2 | 0.04498 |
| 1100 | Q55574 | -0.03864 | -0.1159 | 3 | 0.06126 |
| 1100 | Q55574 | -0.03864 | -0.1545 | 4 | 0.09027 |
| 1100 | Q55626 | -0.005311 | -0.007966 | 1.5 | 0.05902 |
| 1100 | Q55626 | -0.005311 | -0.01062 | 2 | 0.08786 |
| 1100 | Q55643 | -0.009459 | -0.01419 | 1.5 | 0.03819 |
| 1100 | Q55643 | -0.009459 | -0.01892 | 2 | 0.05588 |

| Light condition (μmol photons $m^{-2}$ $s^{-1}$) | Enzyme (UniProt ID) | Lower bound, original (mmol $gDW^{-1}$ $h^{-1}$) | Lower bound, relaxed (mmol $gDW^{-1}$ $h^{-1}$) | Fold-change | μ after relaxation ($h^{-1}$) |
|---|---|---|---|---|---|
| 1100 | Q55643 | -0.009459 | -0.02838 | 3 | 0.07472 |
| 1100 | Q55664 | -4.286 | -214.3 | 50 | 0.00699 |
| 1100 | Q55759 | -0.1616 | -0.2424 | 1.5 | 0.00785 |
| 1100 | Q55759 | -0.1616 | -0.3232 | 2 | 0.0119 |
| 1100 | Q55759 | -0.1616 | -0.4848 | 3 | 0.01539 |
| 1100 | Q55759 | -0.1616 | -0.6465 | 4 | 0.02379 |
| 1100 | Q55759 | -0.1616 | -0.8081 | 5 | 0.03078 |
| 1100 | Q55759 | -0.1616 | -1.616 | 10 | 0.03909 |
| 1100 | Q55759 | -0.1616 | -8.081 | 50 | 0.07825 |
| 1100 | Q57417 | -0.03995 | -0.05993 | 1.5 | 0.05083 |
| 1100 | Q57417 | -0.03995 | -0.07991 | 2 | 0.07576 |
